# No Transcriptional Compensation for Extreme Gene Dosage Imbalance in Fragmented Bacterial Endosymbionts of Cicadas

**DOI:** 10.1101/2023.04.02.535232

**Authors:** Noah Spencer, Piotr Łukasik, Mariah Meyer, Claudio Veloso, John P. McCutcheon

## Abstract

Bacteria that form long-term intracellular associations with host cells lose many genes, a process that often results in tiny, gene-dense, and stable genomes. Paradoxically, the same evolutionary processes that drive genome reduction and simplification may also sometimes cause genome expansion and complexification. A bacterial endosymbiont of cicadas, *Hodgkinia cicadicola*, exemplifies this paradox. In many cicada species, a single *Hodgkinia* lineage with a tiny, gene-dense genome has split into several interdependent cell and genome lineages. Each new *Hodgkinia* lineage encodes a unique subset of the ancestral unsplit genome in a complementary way, such that the collective gene contents of all lineages sum to the total found in the ancestral single genome. This splitting process creates genetically distinct *Hodgkinia* cells that must function together to carry out basic cellular processes, but also creates a gene dosage problem where some genes are encoded by only a small fraction of cells while others are much more abundant. Here, by sequencing DNA and RNA of *Hodgkinia* from different cicada species with different amounts of splitting – along with those of its structurally stable, unsplit partner endosymbiont *Sulcia muelleri* – we show that *Hodgkinia* does not transcriptionally compensate to rescue the wildly unbalanced gene and genome ratios that result from lineage splitting. We also find evidence that *Hodgkinia* has a reduced capacity for basic transcriptional control independent of the splitting process. Collectively, these findings reveal another layer of degeneration that further pushes the limits of canonical molecular and cell biology in *Hodgkinia*, and may in part explain its propensity to go extinct through symbiont replacement.

**Significance:** Many cicadas host two bacterial endosymbionts, *Hodgkinia* and *Sulcia*, which produce essential amino acids missing from the insect’s xylem sap diet. Following 100+ million years of strict host association, both bacteria have lost many genes and posses extremely tiny genomes. In some cicadas, *Hodgkinia* has split into multiple cell lineages, distributing its genes, with little respect to their function, among separate lineages present at (sometimes wildly) different abundances. We find no transcriptional response to this in *Hodgkinia*, resulting in similarly imbalanced mRNA abundances. We also find less control of transcription in *Hodgkinia* compared to *Sulcia*. *Hodgkinia*’s transcriptome embodies an extreme, even relative to other highly-reduced endosymbionts, and raises questions about how cell biology in multi-lineage *Hodgkinia* can function at all.

## Introduction

Vertically transmitted bacterial endosymbionts that form very stable and long-term association with host cells, including the mitochondrion and plastid of eukaryotic cells, can lose most of the genes originally encoded by their free-living ancestors (Andersson and Kurland 1998; Green 2011; Gray 2012). Endosymbiont genomes are often small in size, stable in structure, and densely packed with a core set of functional genes (Boore 1999; Tamas et al. 2002; McCutcheon and Moran 2011; Graf et al. 2021). While such tiny, stable, and gene-dense endosymbiont genomes have evolved again and again in diverse host lineages, some endosymbiont and organelle genomes have secondarily become unstable, expanding in size through the accumulation or proliferation of non-coding and non-functional DNA. The cicada endosymbiont *Candidatus* Hodgkinia cicadicola (hereafter, *Hodgkinia*) and the mitochondria of some sucking lice and flowering plants have all evolved multichromosomal genomes several times larger than those of closely related lineages despite virtually no change to their overall gene repertoire (Shao et al. 2012; Sloan et al. 2012; Campbell et al. 2015; Campbell et al. 2017). In the case of *Hodgkinia*—and in contrast to mitochondria, where the distinct chromosomes are mixed together throughout the mitochondrial compartments of a cell—genome fragmentation occurs in parallel with cellular diversification such that the total gene set is divided among distinct *Hodgkinia* cell populations which are present at different relative abundances in the host (Van Leuven et al. 2014; Łukasik et al. 2018). As a result, genes critical both to *Hodgkinia*’s symbiotic role in nutrient biosynthesis along with genes central to basic bacterial cell function can differ in abundance by orders of magnitude within the same insect. This gene dosage problem raises the question of whether complex *Hodgkinia* can correct for large differences in gene abundance in some way, for example through transcriptional up-regulation of lowly abundant genes (Campbell et al. 2015; Łukasik et al. 2018).

The *Hodgkinia* genome has the expected single circular-mapping chromosome structure in many cicadas (McCutcheon et al. 2009b; Van Leuven et al. 2014; Łukasik et al. 2018). In some cicadas, however, *Hodgkinia* has independently undergone varying degrees of genome fragmentation via cell lineage splitting (Łukasik et al. 2018; Campbell et al. 2017). Compared to the unsplit ancestral genome, individual split genomic lineages lack functional copies of many essential genes, but these losses occur in a complementary fashion such that the unsplit gene set is maintained at the level of the total *Hodgkinia* population in each cicada (Van Leuven et al. 2014; Łukasik et al. 2018). The complementary genome erosion of each lineage enforces transmission of all *Hodgkinia* genomes to the subsequent host generation, resulting in an expansion of the total *Hodgkinia* genome from the perspective of the host (Campbell et al. 2015). In extreme cases, this splitting process results in genome complexes consisting of at least a dozen lineages and totaling over 1.5 Mb in length, a more than tenfold increase in genome size relative to single-lineage *Hodgkinia* genomes (Campbell et al. 2017). Importantly, comparisons between these largest *Hodgkinia* complexes show extreme variation in splitting outcomes with respect to the size and gene content of their constituent genomes, which suggests that splitting does not converge on a particular endpoint or optimum (Campbell et al. 2017).

*Hodgkinia*’s predisposition to splitting may owe in part to its high rate of sequence evolution, a feature also observed in the huge, fragmented mitochondrial genomes of the angiosperms *Silene conica* and *Silene noctiflora* (Sloan et al. 2012; Van Leuven et al. 2014). This is contrasted by the roughly 50–100 times lower nucleotide substitution rate exhibited by *Hodgkinia*’s partner endosymbiont, *Candidatus* Sulcia muelleri (hereafter, *Sulcia*) (Van Leuven et al. 2014). While *Hodgkinia* genomes are structurally unstable and vary widely in size, *Sulcia* tends to be much more stable. Following at least 250 million years of strict host-association, *Sulcia* genomes from distantly related hosts show almost perfect gene co-linearity and very similar gene sets (Moran et al. 2005; Bennett and Moran 2015) [although a broader sampling of Auchenorrhynchan insects shows that several genomic inversions have occurred in different *Sulcia* lineages (Deng et al. 2022)]. Likewise, while several cicada groups have replaced *Hodgkinia* with fungal endosymbionts, *Sulcia* is retained in every cicada species examined to date (Matsuura et al. 2018; Wang et al. 2022).

Given the diversity of *Hodgkinia* genome size and organization and the relative structural stasis of *Sulcia* genomes in cicadas, this system constitutes an elegant natural experiment for evaluating the downstream transcriptional consequences of wild swings in gene dosage resulting from endosymbiont genome instability. To characterize the transcriptional activity of *Hodgkinia* and *Sulcia* genomes relative to their genomic abundance, we sequenced DNA and RNA from the symbiotic organs of 18 cicadas representing 6 species encompassing a spectrum of *Hodgkinia* complexity. We find that *Hodgkinia* exerts limited transcriptional control compared to *Sulcia* and is unable to transcriptionally compensate for the massive effect of gene dosage imbalance that is produced by lineage splitting.

## Results

In the absence of lineage splitting, we assume each *Hodgkinia* cell contributes equally to the total abundance of each *Hodgkinia* transcript (**Figure 1A**). Following splitting and differential gene loss, some transcripts can only be produced by a (sometimes very small) subset of *Hodgkinia* cells. We evaluated four different hypotheses, two adaptive and two non-adaptive, for how *Hodgkinia* may or may not compensate at the transcriptional level for the gene dosage imbalances that result from splitting and differential gene loss: an adaptive response of non-specific, constitutive transcriptional upregulation to bring transcripts of low-abundance genes to some threshold level (“overcompensation,” **Figure 1B**); a specific adaptive response where lowly abundant genes are upregulated to rescue pre-splitting transcript abundances (“complementation,” **Figure 1C**); a non-response, where each gene is transcribed at its pre-splitting levels in each cell irrespective of its relative abundance (“subdivision,” Figure 1C); and a response of further regulatory decay, where non-compensatory changes to transcription are introduced as a side-effect of splitting and genome erosion (“disruption,” **Figure 1D**). To look for signatures of these outcomes across the spectrum of *Hodgkinia* complexity, we collected three individuals from single populations representing each of six different cicada species (**Table 1)** and sequenced the metagenomes and metatranscriptomes of their dissected bacteriomes (endosymbiont-housing organs).

**Figure 1:**
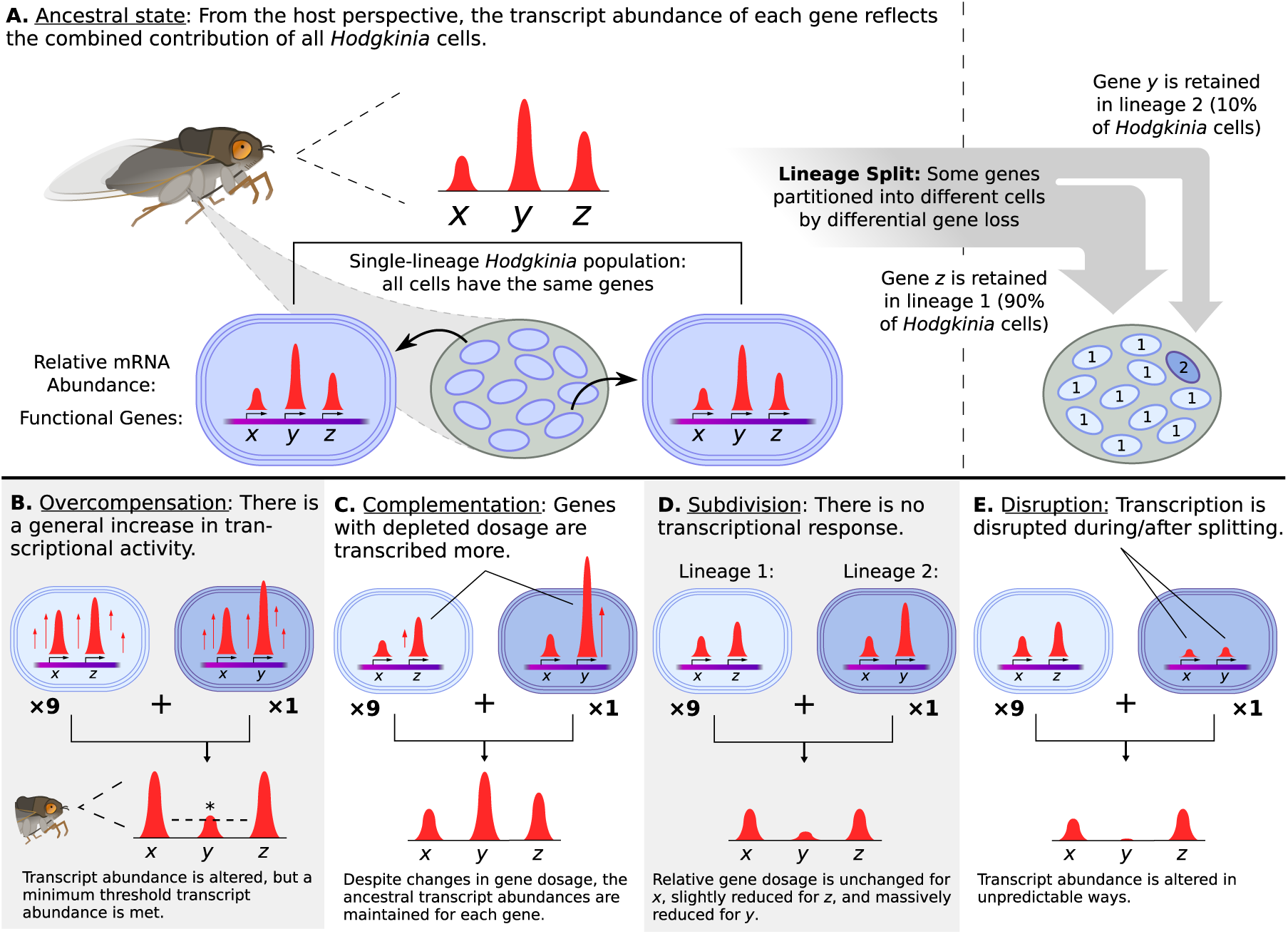
Schematic of a Hodgkinia cell lineage split and its possible transcriptional outcomes. **(A)** In the absence of cell lineage splitting, Hodgkinia cells all contain the same genes (here x, y, and z) and contribute to their respective transcript abundances. Lineage splitting and complementary gene loss decrease the relative dosage (total supply) of genes which are lost in some cells. In this abstracted example, genes x, y, and z have relative post-splitting dosages of 1.0 (full dosage), 0.1, and 0.9, respectively. **(B)** If Hodgkinia cells increase transcription genome-wide, transcript abundances will remain imbalanced but could reach some required threshold level for dosage-depleted genes. **(C)** If Hodgkinia transcription is regulated to transcribe dosage-depleted genes at higher levels, pre-splitting transcript abundances could be rescued. **(D)** If Hodgkinia cells do not change transcription in response to changes in gene dosage (i.e. each gene is transcribed at roughly its original level in each cell), transcript abundance of genes with reduced dosage will decrease. **(E)** If the processes of cell lineage splitting and/or gene loss intrinsically affect transcription of certain genes, transcript abundances could change in unpredictable ways.

**Table 1.**
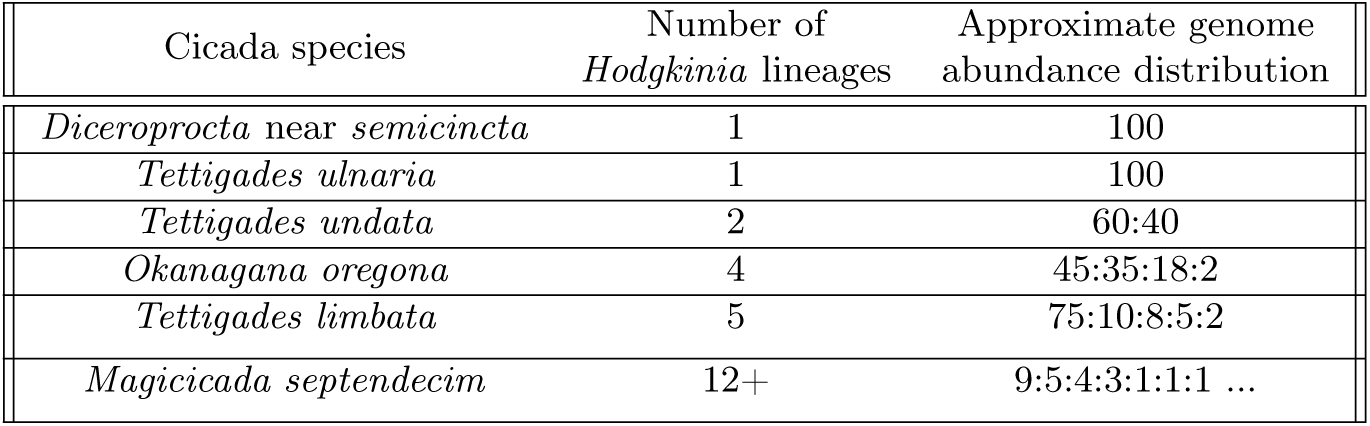
Hodgkinia genome complexity and abundance ratios in all cicada species sampled.

The multi-lineage *Hodgkinia* studied here all originated from independent splitting events (Camp-bell et al. 2017; Łukasik et al. 2018). Closed genomes are available for all relevant *Hodgkinia* lineages except in *M. septendecim*, in which the *Hodgkinia* genome has been assembled into 39 circular molecules and 124 additional contigs. Similarly, closed genomes of all relevant *Sulcia* lineages are available, with the exception of *Sulcia* from *M. septendecim*. In this case, we used the genome of *Sulcia* from *Magicicada tredecim*, which is completely co-linear with and over 99% identical to its counterpart in *M. septendecim*. We obtained between 18.7 and 44.4 million paired-end reads from the bacteriome metagenome libraries and between 43.6 and 181.7 million reads from the corresponding metatranscriptome libraries. Each of these libraries contain sequences derived from *Hodgkinia*, *Sulcia*, and the cicada host.

### Cicada Endosymbionts Retain Different Degrees of Transcriptional Control

We began by examining transcription in the cicada endosymbionts in general. Relatively little work has characterized transcription in endosymbionts with extremely reduced genomes (Bennett and Chong 2017; Van Leuven et al. 2019; Wang et al. 2022), but a comparative analysis of RNA polymerase genes suggests that some endosymbionts, including *Hodgkinia*, may have a limited capacity for promoter recognition (Rangel-Chávez et al. 2021). To compare the degree to which *Hodgkinia* and *Sulcia* can specifically transcribe coding DNA and can transcribe genes at different levels in line with biological expectations, we aligned stranded bacteriome mRNA-seq reads from each cicada species to the corresponding *Sulcia* (**Figure 2A–B;** Supplementary Figure S1) and *Hodgkinia* (**Figure 2C–D; Supplementary Figures S2–S5**) reference genomes and visualized per-base coverage across each chromosome. We obtained >1X coverage of the vast majority of genomic regions, except for some of the small, unplaced *Hodgkinia* contigs in *M. septendecim*. In both endosymbionts and across host species, patterns of coverage were highly consistent among biological replicates (**Figure 2; Supplementary figures S1–S5**). In the case of *Sulcia*, we saw clear similarities between species in the relative transcription of different genes (**Figure 2A–B; Supplementary Figure S1**), while *Hodgkinia* was much more variable (**Figure 2C–D; Supplementary Figures S2–S5**).

**Figure 2:**
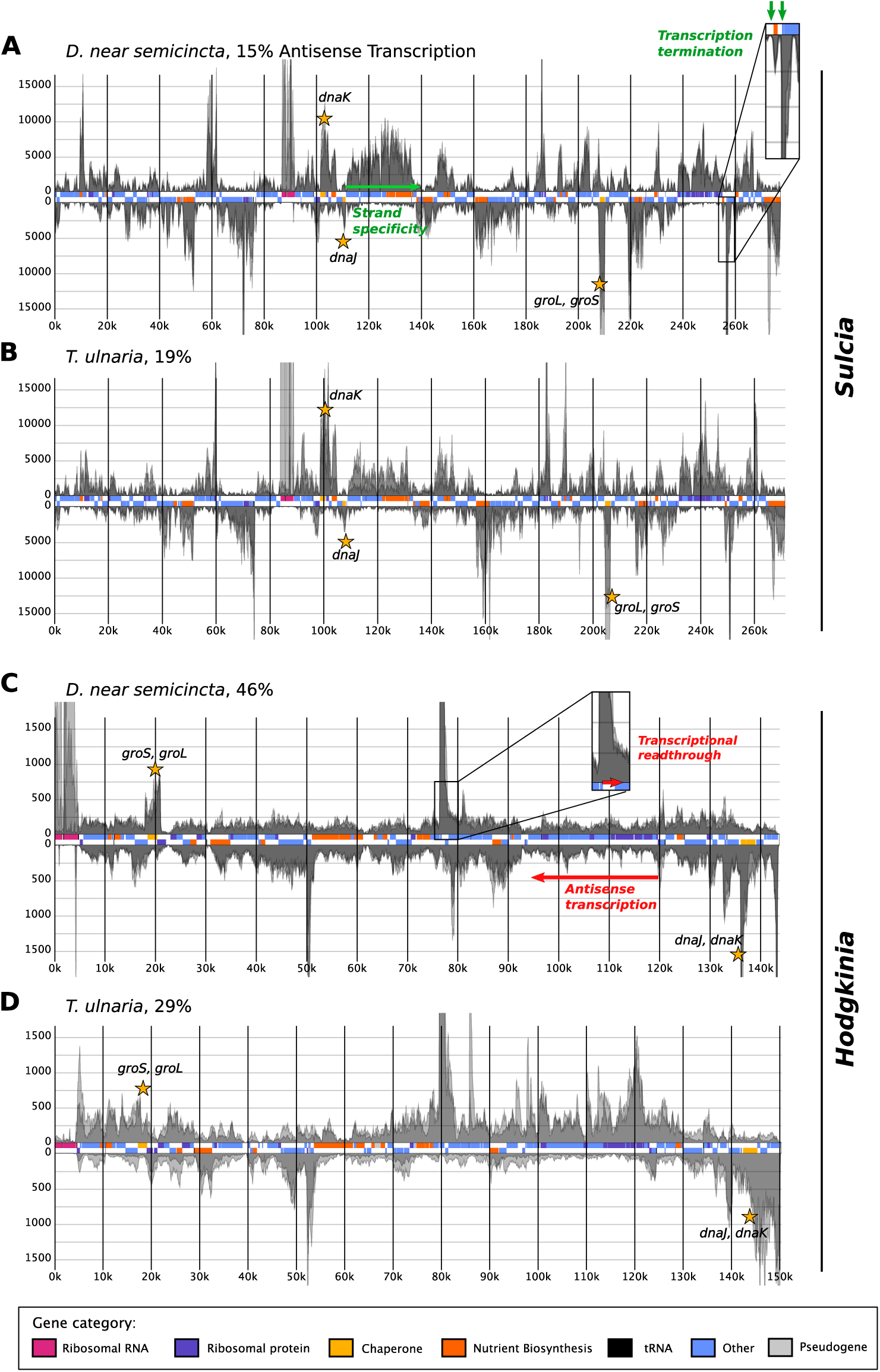
Strand-specific, per-base RNA-seq coverage along the chromosomes of Sulcia (A–B) and Hodgkinia (C–D) from D. near semicincta and T. ulnaria. Rectangles in the central track of each plot represent annotated genes and are colored according to functional categories. Positive and negative Y axes correspond to coverage of unfiltered RNA-seq reads derived from the plus and minus strands of each chromosome, respectively. For each plot, coverage profiles from each biological replicate (translucent gray) are overlaid along the same axes. Panels A and B represent alignments downsampled to approximately 3500X mean coverage of the Sulcia genome and are cropped at approximately y=±20000. Panels C and D represent alignments downsampled to approximately 450X mean coverage of the Hodgkinia genome and and are cropped at approximately y=±2000. Antisense counts as a percentage of sense+antisense counts are shown for each genome.

**Figure 5:**
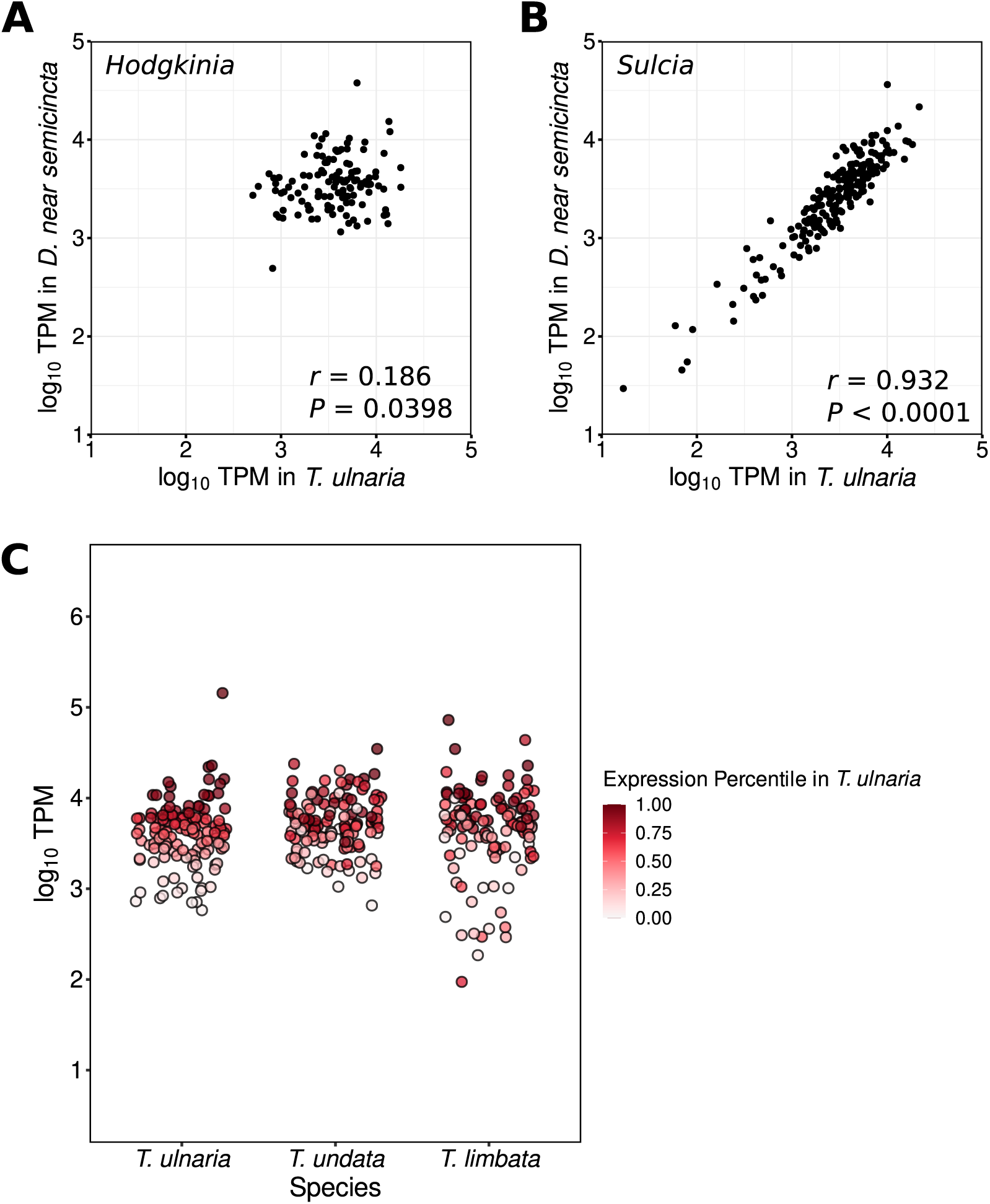
Differences in homologous gene expression between endosymbionts of different host species. Scatter plots show the correlation of relative expression levels (log_10_ TPM) for homologous protein-coding genes in two symbionts, **(A)** single-lineage Hodgkinia and **(B)** Sulcia, between two distantly related cicadas: D. near semicincta and T. ulnaria. The Pearson correlation coefficient r and the p-value for a test of correlation are given. **(C)** Expression levels of homologous Hodgkinia genes in T. ulnaria, T. undata, and T. limbata are shown colored according to their TPM expression percentile in T. ulnaria, highlighting variation in Hodgkinia transcript abundances within the genus Tettigades. All TPM values are averages across biological replicates. TPM values in T. undata and T. limbata represent summed values from all copies of a given gene.

Compared with *Hodgkinia*, *Sulcia* exhibited patterns of transcription which indicate a greater ability to terminate transcription. *Sulcia* exhibited clearly distinguishable peaks of high RNA coverage overlapping with annotated genes (inset of **Figure 2A**). In *Hodgkinia*, RNA coverage was often (but less consistently) high along annotated genes. However, rather than producing symmetrical peaks of coverage centered on annotated genes, transcription in *Hodgkinia* frequently continued past the ends of genes, gradually decreasing until another peak of RNA coverage began. For example, the RNase P RNA gene is transcribed at high levels in *Hodgkinia* from both *D.* near *semicincta* and *T. ulnaria* (**Figure 2C–D**), but the corresponding peak in RNA-seq coverage continues for several times the length of the functional RNA and into an intergenic region (inset of **Figure 2C**). Unlike in protein-coding genes, several of which show similar “run-on” transcriptional profiles in *Hodgkinia*, the transcriptional readthrough at this locus cannot be explained or resolved by termination at the level of translation.

The *Hodgkinia* and *Sulcia* lineages sampled often appeared to highly transcribe chaperone genes (*groS*, *groL*, *dnaJ*, and *dnaK* ; **Figure 2**, gold stars). This is consistent with published endosymbiont transcriptomes (Stoll et al. 2009; Luck et al. 2015; Medina Munoz et al. 2017) and with proteomic data showing that chaperones are among the most abundant proteins in endosymbionts (Charles et al. 1997; McCutcheon et al. 2009a; McCutcheon et al. 2009b; Poliakov et al. 2011).

To evaluate transcriptional control more quantitatively, we calculated levels of antisense transcription in *Hodgkinia* and *Sulcia* from each cicada species. To do this, we first used the transcript quantification tool FADU to obtain transcript counts for functional genes in *Sulcia* and *Hodgkinia* (excluding tRNA and rRNA genes) (Chung et al. 2021). We then repeated this step for antisense transcription by deliberately specifing the opposite strand orientation for our libraries such that FADU output counted alignments to the strand opposite each open reading frame (Srinivasan et al. 2020). *Hodgkinia* had a higher proportion of antisense counts than its coresident *Sulcia* in every biological replicate from each cicada species (**Supplementary Figures S1–S5**). *Hodgkinia* from *D*. near *semicincta*, which has experienced no genome fragmentation, stood out in this regard with an average of 46% antisense transcripts compared to just 15% in its coresident *Sulcia* (**Figure 2A, 2C**).

Taken together, these data show an overall loss of transcriptional control in *Hodgkinia* compared to *Sulcia* (**Supplementary Figures S1–S5**), and provide the first indirect hint that dosage compensation at the level of *Hodgkinia* transcription is unlikely to be occurring in these symbioses.

### Complex Hodgkinia Produce RNA in Proportion to Their Cell Abundance

Under our first hypothesized adaptive scenario, complex *Hodgkinia* could rescue the transcript abundances of low-copy genes through a general increase in mRNA synthesis (overcompensation, **Figure 1B**). We determined the relative contributions of *Hodgkinia* and *Sulcia*-derived DNA and mRNA to the sequencing libraries from each specimen by filtering out any remaining ribosomal RNA sequences, mapping each set of filtered reads to the corresponding endosymbiont genomes, and calculating the coverage of each genome as a proportion of all filtered reads (**Figure 3**). We have already shown that *Hodgkinia* genome coverage is a good proxy for cell abundance (Van Leuven et al. 2014; Campbell et al. 2018; Łukasik et al. 2018). The DNA abundance of *Sulcia* was consistently higher than that of *Hodgkinia* except in the *M. septendecim* samples, which also had the highest overall *Hodgkinia* DNA abundance.

**Figure 3:**
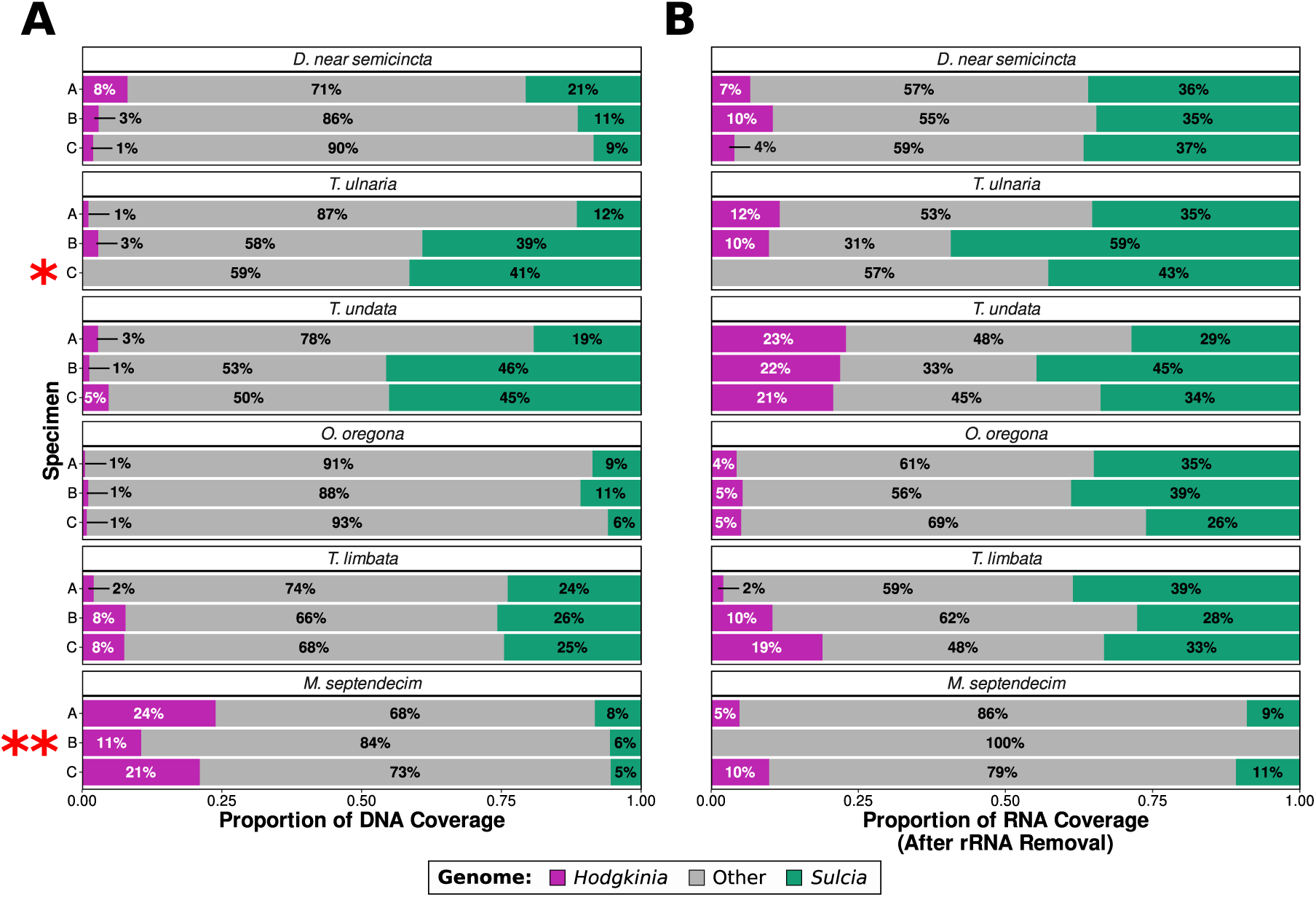
Proportional contributions of Hodgkinia and Sulcia to the total (A) DNA and (B) RNA sequencing coverage, shown for triplicate biological replicates of each cicada species examined. Reads mapping to the Hodgkinia and Sulcia rRNA genes for each cicada were removed prior to calculation of RNA coverage. T. ulnaria specimen C (one asterisk) lacked Hodgkinia DNA and RNA and was excluded from subsequent Hodgkinia-based analyses. M. septendecim specimen B (two asterisks) lacked RNA coverage from either endosymbiont and was excluded from all further analyses.

Compared to the DNA libraries, the RNA libraries generally contained more endosymbiont-derived reads. In an overcompensation scenario, complex *Hodgkinia* would be expected to produce a greater ratio of RNA:DNA coverage than their coresident *Sulcia*. While *Hodgkinia* from *T. un-data* showed patterns of coverage potentially consistent with overcompensation in all three biological replicates (for example, specimen A showed 3% *Hodgkinia* coverage in DNA reads but 23% coverage in RNA reads), we did not observe this pattern in any of the other multi-lineage *Hodgkinia* examined. In the samples representing the most extreme level of splitting, *Hodgkinia* from *M. septendecim*, where overcompensation might be expected to be the most obvious, RNA coverage was actually underrepresented relative to its DNA abundance (for example, specimen C showed 21% *Hodgkinia* coverage in DNA reads but only 10% coverage in RNA reads). This decrease in relative RNA abundance was not an artifact of rRNA depletion being less effective in certain RNA-seq libraries, as the trends we observed in total RNA coverage fractions hold even when rRNA is not removed bioinformatically (**Supplementary Figure S6**).

One sample from *T. ulnaria* contained very little *Hodgkinia* material and was excluded from any other *Hodgkinia*-based analysis (marked with an asterisk in **Figure 3**). A sample from *M. septendecim* produced very little endosymbiont-derived RNA coverage in general, with *Hodgkinia* and *Sulcia* collectively contributing less than 0.1%, and was excluded from all other analysis (marked with two asterisks in **Figure 3**). Because cicadas live underground for most of their lives, only emerge once a year (at most), and are difficult to catch, we were unable to add new samples to replace these lost data points.

### Gene Dosage Depletion Reshapes the Hodgkinia Transcriptome

Under a complementation scenario (**Figure 1C**), the distribution of transcript abundances in the total *Hodgkinia* population (i.e. from the perspective of the insect host) would be similar between single-lineage and complex *Hodgkinia*. Conversely, in the absence of complementation, genes which are present in fewer copies in the *Hodgkinia* population following splitting would be represented by fewer transcripts than more abundant genes. To distinguish these two outcomes, we compared relative gene dosage with total transcript abundance in each *Hodgkinia* system. In each biological replicate, we defined the dosage or abundance of a gene as the percentage of *Hodgkinia* DNA sequencing coverage contributed by *Hodgkinia* contigs which contain the gene (**Supplementary Table S1**). Similarly, we measured total transcript abundance of a gene as the summed transcripts per million (TPM) of each distinct copy of the gene in a *Hodgkinia* complex (Li and Dewey 2011; Wagner et al. 2012).

The TPM distributions from single-lineage *Hodgkinia* systems (*D.* near *semicincta* and *T. ulnaria*, **Figure 4A–B**) showed relatively consistent shapes among biological replicates but differed slightly in spread between the two host species, comparable to the between host variation in *Sulcia* TPM distributions (**Figure 4G–L**). The TPM distributions from *T. undata*, hosting two *Hodgkinia* lineages, were similar in shape but showed a clear bi-forcation on the basis of gene dosage with genes at full dosage corresponding to upper half of the distributions and genes at 40-60% relative dosage corresponding to the lower half (**Figure 4C**). Compared to these three species, the TPM distributions from the more fragmented *Hodgkinia* of *O. oregona* and *T. limbata* showed relatively more genes at their extreme low ends (**Figure 4D–E**), and most of these genes had relative dosages of less than 20% in these cicadas. This was not the case in the highly-fragmented *Hodgkinia* of *M. septendecim* in which all genes are far from maximal dosage. Instead, its TPM distributions showed uniquely high dispersion with relatively few values concentrated around the average (**Figure 4F**). Such differences in the shape of the TPM distribution were specific to *Hodgkinia*: they were not observed in their coresident *Sulcia* lineages (**Figure 4G–L**).

**Figure 4:**
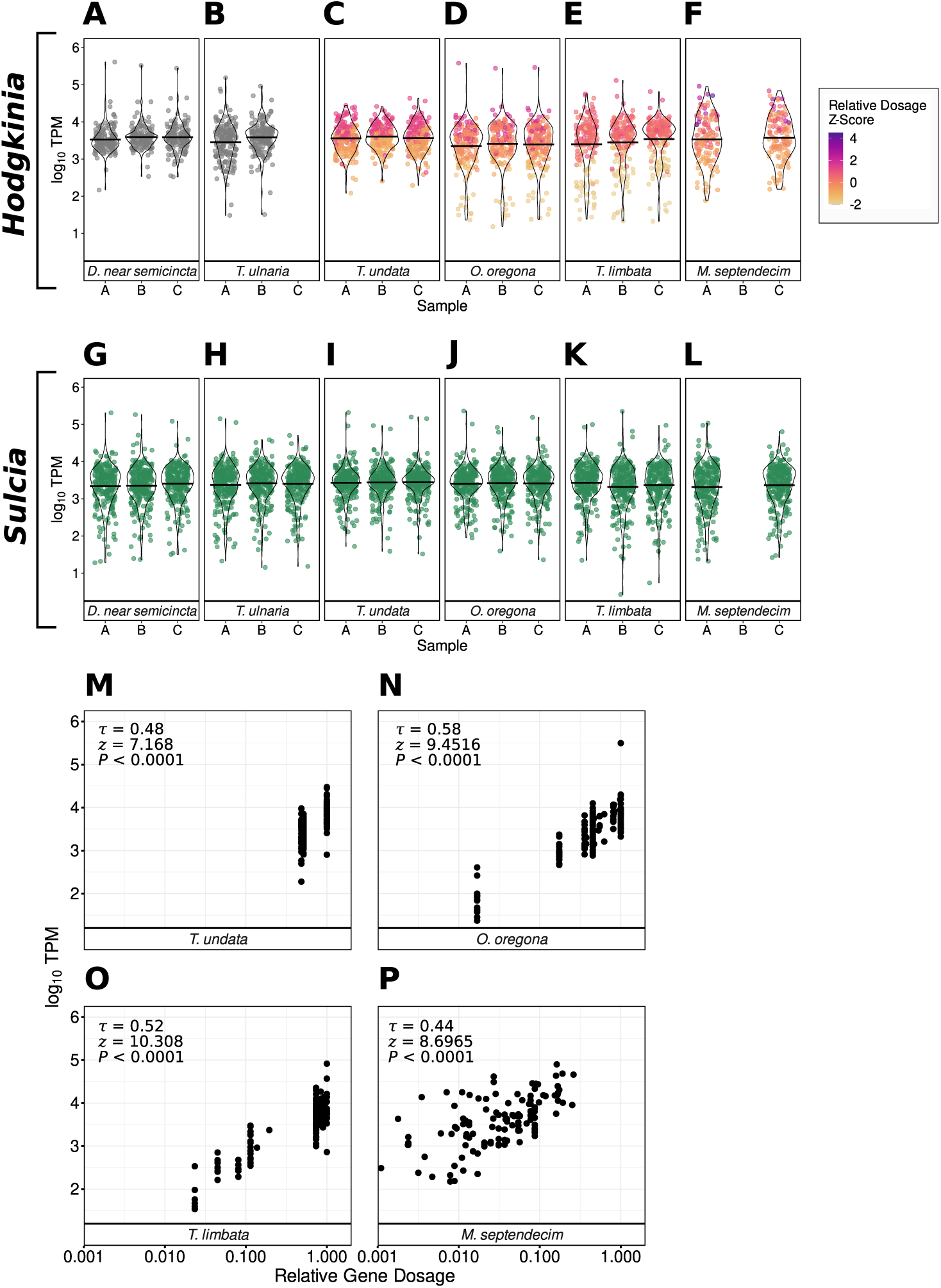
Endosymbiont gene dosage and gene expression from the host perspective in symbiotic systems of varying complexity. **(A–F)** Distribution of log_10_ TPM expression levels of Hodgkinia genes from each cicada specimen examined. Points represent the summed TPM of all copies of a gene. Points are colored according to their relative dosage. To make differences in gene dosage visible in M. septendecim, gene dosages for all specimens were converted to standard deviations above or below the specimen’s average Hodgkinia gene dosage (Z transformation). **(G–L)** Distribution of log_10_ TPM expression levels of Sulcia genes from each cicada specimen examined. **(M–P)** Total log_10_ TPM expression levels (y-axis) of Hodgkinia genes of different relative dosage (x-axis, log_10_ scale) in Hodgkinia systems of varying complexity. TPM and relative dosage values are averages of values from each biological replicate. For each comparison, Kendall’s rank correlation coefficient τ as well as the test statistic (Z) and p-value for hypothesis tests of rank correlation are given, showing a significant positive correlation between relative Hodgkinia gene dosage and transcript abundance in all four species.

In all cicada species with split *Hodgkinia*, we found a significant positive correlation between relative gene dosage and total TPM at a significance threshold of *α* = 0.05 (Kendall’s rank correlation, P < 0.0001 for all four comparisons, **Figure 4M–P**). In other words, *Hodgkinia* genes which had greater dosage at the genomic level in each cicada tended to be represented by a greater number of transcripts. The strength of the relationship, given by Kendall’s *τ* , was similar across host species, ranging from 0.44 in *M. septendecim* to 0.58 in *O. oregona*.

This correspondence between total gene and transcript abundances in multi-lineage *Hodgkinia* is inconsistent with complementation. However, complementary changes in transcription could still exist, even if they don’t overcome the influence of gene dosage altogether. We tested for cell-level complementary responses to differences in total gene supply using semi-partial Kendall rank correlations. Semipartial correlations allowed us to characterize the relationship between the TPM abundance of all distinct *Hodgkinia* gene copies and their relative dosage from the host perspective while controlling for the effect of each gene copy’s DNA abundance on its measured transcript abundance. Per-cell transcription was expected to be negatively correlated with relative gene dosage under a complementation scenario (**Figure 1C**) and not correlated under a subdivision scenario (**Figure 1D**). We found no significant correlation between cell abundance-controlled TPM and relative gene dosage in *O. oregona* (*τ* = 0.018, *Z* = 0.44, P = 0.66), in *T. limbata* (*τ* = 0.053, *Z* = 1.372, P = 0.17), or in *M. septendecim* (*τ* = 0.054, *Z* = 1.268, P = 0.205) at a significance threshold of *α* = 0.05. In *T. undata*, abundance-controlled TPM and relative gene dosage were significantly positively correlated, indicating that genes encoded in only one of *T. undata*’s two *Hodgkinia* cell lineages actually tended to have lower per-cell expression (*τ* = 0.209, Z = 5.071, P < 0.0001).

### Hodgkinia Transcription Profiles Are Not Conserved Across Host Species

Having found no evidence for transcriptional compensation for the gene dosage outcomes resulting from *Hodgkinia* lineage splitting and reciprocal gene loss, we next asked whether gene expression patterns in the non-fragmented ancestral *Hodgkinia* transcriptome are conserved between species. We compared the log_10_ TPM expression of homologous, protein-coding *Hodgkinia* and *Sulcia* genes between *D*. near *semicincta* and *T. ulnaria*, which both host a single *Hodgkinia* lineage (**Figure 5A–B**). Expression of Sulcia genes showed a strong linear correlation between the two host species (Pearson correlation: r = 0.932, t = 36.275, *ν* = 198, P < 0.0001) while *Hodgkinia* gene expression was only weakly correlated (r = 0.186, t = 2.087, *ν* = 121, P = 0.039). In addition to the strong biological contrast between these outcomes, the consistency of *Sulcia* transcriptional profiles across relatively distantly related host species gives us confidence that our RNA-seq data are of good overall quality and that the relatively noisy nature of the *Hodgkinia* data is not the result of technical artifacts.

Given the incongruence between transcript abundances in phylogenetically distant single lineage *Hodgkinia*, we directed our focus to the genus *Tettigades*, from which we had sampled three different species, reasoning that *Hodgkinia* transcript abundances in *T. ulnaria* (which hosts a single *Hodgkinia* lineage) may approximate a pre-splitting “starting point” for this group, and that some semblance of transcriptional control may persist in phylogenetically related lineages. Total TPM transcript abundances in multi-lineage *Hodgkinia* from *Tettigades* cicadas appear to deviate from this hypothetical starting point, even in the case of *T. undata*, which hosts only two different *Hodgkinia* lineages (**Figure 5C**).

## Discussion

### Hodgkinia Does Not Compensate for Transcriptional Consequences of Gene Dosage Imbalance

The process of splitting into multiple interdependent cell lineages combined with complementary gene loss has resulted in varied and sometimes extreme gene dosage outcomes for the *Hodgkinia* populations contained in each cicada (Campbell et al. 2017; Łukasik et al. 2018). We considered two possible outcomes that would reflect compensation for this change at the level of transcription: widespread overproduction of mRNA to guarantee sufficient transcript abundance (overcompensation, **Figure 1B**) and fine-tuned compensatory regulation to rescue the transcription levels of dosage-depleted genes (complementation, **Figure 1C**). Our analysis of genome relative abundance and transcription in *Hodgkinia* of multiple complexity levels shows that neither of these adaptive responses occur. Rather, the transcriptional changes that occur in *Hodgkinia* composed of 2, 4, 5, and 12+ cell lineages consistently, strongly, and simply reflect the gene dosage of corresponding genes on their genomes.

Some form of compensation could, in principle, occur at the level of translation. This would presumably rely on factors external to *Hodgkinia*, particularly since the least abundant *Hodgkinia* cell lineages in the species examined here tend to encode relatively limited complements of translation-related genes compared to more abundant lineages (Campbell et al. 2017; Łukasik et al. 2018). Our observations could also be affected by the age of the cicadas sampled, which, as fully grown adults nearing the ends of their lives, may no longer be as reliant on the proper functioning of their nutritional endosymbionts in one or both sexes. However, given the lack of conservation in *Hodgkinia* transcript abundance across cicada species, and the relative conservation we see in *Sulcia* transcription, we favor the idea that *Hodgkinia* simply tolerates the transcriptional consequences of gene dosage changes, even quite extreme ones.

This is not to say that such changes are always selectively neutral in *Hodgkinia*. Given that our results are most consistent with a lack of transcriptional response in *Hodgkinia* after splitting, our finding that genes which had been lost in one of *T. undata*’s two *Hodgkinia* lineages had lower per-cell transcription could suggest that lineage-specific gene losses are more likely to be fixed when they occur in lowly-expressed genes. Additionally, while the dosage outcomes in highly complex *Hodgkinia* are not deterministic, some mechanism seems to favor the retention of certain *Hodgkinia* genes at a greater total abundance, and this is reflected in their transcript abundance (Campbell et al. 2017; Łukasik et al. 2018).

### Basic Transcriptional Control Shows Signs of Erosion in Hodgkinia

The transcriptional machinery encoded by the bacterial endosymbionts with the tiniest genomes is extremely rudimentary (McCutcheon and Moran 2011). Despite this, we found evidence for at least some degree of transcriptional control in two such endosymbionts. All *Sulcia* and *Hodgkinia* transcriptomes produced more alignments to genes in the sense orientation than in the antisense orientation, suggesting that open reading frames are preferably transcribed over random positions on the opposite strand of DNA. We also observed consistently high chaperone gene expression in both endosymbionts, a recurring feature of endosymbiont transcriptomes thought to be of functional importance to the endosymbiotic lifestyle (Fares et al. 2002; Stoll et al. 2009; McCutcheon and Moran 2011; Luck et al. 2015; Medina Munoz et al. 2017). This occurs despite the loss of the *rpoH* -encoded *σ*^32^ heat shock sigma factor, which modulates expression of chaperone genes in free-living bacteria (Neidhardt and VanBogelen 1981; Yamamori and Yura 1982; Grossman et al. 1987).

Across all six host species examined, *Hodgkinia* and *Sulcia* differed in two potential indicators of transcriptional control. First, we found that RNA-seq coverage declined predictably at gene ends in *Sulcia* while the high coverage typical of transcriptional start sites frequently extended past annotated genes in *Hodgkinia*, possibly indicating transcriptional read-through. The gene contents of these two endosymbionts point to a potential mechanistic explanation: *Sulcia*, unlike *Hodgkinia*, retains *rho* and its cofactor *nusA*, which have well-characterized roles in transcription termination (Schmidt and Chamberlin 1984; Richardson 2002; McCutcheon et al. 2009a; McCutcheon et al. 2009b; Łukasik et al. 2018). Second, *Hodgkinia* endosymbionts showed consistently higher levels of antisense transcription than their co-resident *Sulcia*, although this may simply be a reflection of increased transcriptional read-through at genes located adjacent to a gene on the opposite strand.

Surprisingly, the single-lineage *Hodgkinia* endosymbiont of *D*. near *semicincta* stood out in its apparent loss of transcriptional control, exhibiting a considerably higher proportion of antisense transcription than any other endosymbiont lineage we examined. The fact that antisense transcription in *Sulcia* from the *D*. near *semicincta* samples was not correspondingly high suggests that this effect is not a technical artifact. A previous comparative genomic analysis of endosymbiont RNA polymerases identified a deletion seven amino acid residues long in the *σ*_3_ subunit from Hodgkinia in a very closely related cicada species, *Diceroprocta semicincta*, and predicted that this loss could impede recognition of an extended -10 box promoter element (Rangel-Chávez et al. 2021). Promoter elements have not been characterized in *Hodgkinia* or in other endosymbionts with tiny genomes, and we have similarly found no recognizable sequence motifs upstream of *Hodgkinia* or *Sulcia* start codons regardless of transcription level (**Supplementary Figures S7–S8**). However, we note that the *rpoD* gene in *Hodgkinia* from *D*. near *semicincta*, like *D. semicincta*, lacks this portion of the *σ*_3_ subunit found in most other *Hodgkinia* genomes (**Supplementary Figure S9**), although similar deletions in *rpoD* genes in *Hodgkinia* from *M. septendecim* are evidently not accompanied by a correspondingly high level of antisense transcription (**Supplementary Figures S5, S9**).

We also found that *Hodgkinia* transcript abundances in *D*. near *semicincta* were weakly correlated with their homologs’ relative abundances in the single-lineage *Hodgkinia* of *T. ulnaria* in contrast to the strong correlation observed in the *Sulcia* transcriptomes of those cicadas. It is unclear to what extent host-specific losses in *Hodgkinia* transcriptional control may have contributed to this lack of conservation versus the 50+ million years of evolutionary divergence between these *Hodgkinia* lineages (Marshall et al. 2018; Wang et al. 2022).

### Gene Products May Be Spread Extremely Thin in Complex Hodgkinia

On one hand, the unresponsiveness of *Hodgkinia* transcription to extreme gene dosage outcomes is unsurprising given that *Hodgkinia* encodes no transcription factors or alternative sigma factors and has even accumulated functionally important losses to basic transcriptional machinery (McCutcheon et al. 2009b; Galán-Vásquez et al. 2016; Łukasik et al. 2018; Rangel-Chávez et al. 2021). Even in unsplit *Hodgkinia* lineages with uniform gene dosage, precise ratios of relative transcript abundance do not appear to be conserved. On the other hand, *Hodgkinia*’s unresponsiveness is surprising because of what it implies about its biology, specifically its apparent tolerance for extreme unbalancing of essential transcripts’ absolute abundance. In *M. septendecim*, where many genes may be present in fewer than ten percent of cells, we found no evidence for a generalized upregulation of *Hodgkinia* transcription. In fact, *Hodgkinia*’s relative contributions to DNA and RNA coverage in this system imply an overall reduced transcriptional activity.

While *in situ* hybridization has shown that rRNA and genomic DNA are not shared among cells in complex *Hodgkinia*, the endosymbiont’s continued existence necessarily implies the movement of either mRNA, protein, metabolites, or some combination of these between cells by an unknown mechanism (Campbell et al. 2015; Łukasik et al. 2018). The likelihood of a biologically important encounter between two *Hodgkinia* proteins could therefore be limited not just by the abundance of the genes by which they are encoded but also by those genes’ spatial distribution within the cicada bacteriome. In the absence of massive complementation or over-compensation at the level of protein synthesis, it is conceivable that the biochemistry of the most complex *Hodgkinia* occurs slowly or inefficiently relative to their single-lineage counterparts.

### Conclusions

The transcriptomes of cicadas’ bacterial endosymbionts, like their genomes, embody two opposite extremes. *Sulcia* exhibits highly conserved transcript abundance ratios and patterns of RNA-seq coverage that line up with biological expectations. *Hodgkinia*, meanwhile, shows diminished transcriptional control and transcribes genes in proportion to their sometimes wildly imbalanced DNA abundance. In either case, it is difficult to quantify the fitness consequences of these transcriptional outcomes. We expect that at least some of the outcomes we observe in *Hodgkinia*, such as widespread antisense transcription in *D*. near *semicincta* and failure to compensate for massive gene dilution in *M. septendecim*, are costly. The magnitudes of these costs are dependent on translational compensatory changes—if any occur—and, in the latter case, gene product transport. Both of these processes have yet to be characterized in *Hodgkinia*. As with the reproductive burden cicadas experience in order to transmit a complete *Hodgkinia* gene complement to their eggs following extensive lineage splitting, we speculate that these events are costly for the symbiosis and may tip the scales in favor of *Hodgkinia* extinction through symbiont replacement (Campbell et al. 2018; Matsuura et al. 2018; Wang et al. 2022).

## Materials and Methods

### Insect Collection

Adult cicadas, a mixture of males and females, were collected in their natural habitat using insect nets and dissected in the field, with abdomens torn open and placed in 7mL tubes with RNAlater. They were kept refrigerated initially, and, after arrival in the laboratory, stored at -80C until processing.

We preliminarily identified specimens based on morphological characters, and later confirmed identifications using marker gene sequences. In the case of *Diceroprocta*, we collected multiple individuals that we could not distinguish based on morphology, but that represented two genotypes divergent by about 3% within the mitochondrial COI gene, one of which matched the previously characterized *D. semicincta* (Van Leuven and McCutcheon, 2012). Since all individuals represented the other COI genotype, we decided to refer to them as *Diceroprocta* near *semicincta*.

An additional specimen of *Okanagana oregona* collected previously was used for assembly of its respective *Hodgkinia* and *Sulcia* genomes. This specimen was collected and dissected in the same manner, placed in 90% EtOH, and stored at -20C until processing.

#### Collection details

***D.* near *semicincta*** (two males + female) University of Arizona campus, Tucson, AZ, USA, 32.23, -110.95, July 2017

***Tettigades ulnaria*** (three males) Side of the road near Putaendo, Valparaíso Region, Chile, - 32.588, -70.715, Jan 2017

***Tettigades undata*** (three males) Side of the road to Termas de Chillan, Bio Bio Region, Chile, -36.903, -71.537, 6 Jan 2017

***Okanagana oregona*** (three males) Mt. Sentinel, Missoula, MT, USA, 46.86, -113.98, 30 Jun 2017

***Okanagana oregona*** (one male specimen used for *Hodgkinia* and *Sulcia* genome assemblies) Mt. Sentinel, Missoula, MT, USA, 46.86, -113.98, 13 June 2016

***Tettigades limbata*** (male + two females) Hills South of Sierra de Bellavista, O’Higgins Region, Chile, -34.826, -70.742, 13 Dec 2014

***Magicicada septendecim*** (three females) Washington, PA, USA, 40.171, -80.221, 2017

### DNA and RNA Extraction, Library Preparation, and Sequencing

DNA was extracted from carefully dissected bacteriome tissue using the Qiagen DNeasy Blood and Tissue kit (Hilden, Germany) except in the case of the *M. septendecim* samples. For these samples, DNA libraries were prepared from co-extracted DNA obtained during RNA isolation (see RNA work details below). Illumina libraries for all samples were prepared using the Illumina Truseq PCR-free kit (San Diego, CA, USA).

RNA was also extracted from each bacteriome tissue sample using the Qiagen RNeasy Mini kit (Hilden, Germany) according to the included protocol for animal tissue and then DNase-treated using the Invitrogen TURBO DNA-free kit (Waltham, MA, USA). Ribosomal RNA was depleted using the Ribo-Zero Epidemiology Kit from Illumina (San Diego, CA, USA) followed by cleanup with the RNeasy MinElute Cleanup Kit (Hilden, Germany).

The DNA and RNA libraries were sequenced in four batches across a total of six lanes on a HiSeq X instrument in 2x150bp mode at Novogene (Sacramento, CA, USA). One additional DNA library from an *O. oregona* specimen used for genome assembly of its endosymbionts was sequenced on a MiSeq v3 instrument in 2x300bp mode.

### Endosymbiont Genome Assemblies and Annotation

The genomes of *Sulcia* from *T. ulnaria* and *T. limbata* were assembled from previously published cicada bacteriome metagenomes deposited under BioProject accessions PRJNA246493 and PRJNA385844 (Van Leuven et al. 2014; Łukasik et al. 2018). These, as well as *Sulcia* and *Hodgkinia* genomes from *D.* near *semicincta* and *O. oregona* were assembled as follows: reads were trimmed of low-quality ends and adapters using Trimmomatic or Trim Galore! (Bolger et al., 2014) and then merged using Pear (Zhang et al. 2014) or bbmerge (Bushnell et al. 2017). Bacteriome metagenomes were initially assembled using custom installations of SPAdes 3.7.1, 3.11.0, or 3.12.0 (Prjibelski et al. 2020) which were compiled with an increased k-mer length limit of 249 bp. Scaffolds from this assembly were used for blastx searches against a custom database comprising the six frame-translated genomes of several *Hodgkinia* lineages and protein-coding genes from *Sulcia*, twelve other insect-associated and free-living bacteria, cicada mitochondria, and the planthopper *Nilaparvata lugens*. Quality-filtered reads were then re-mapped to scaffolds with top matches (evalue < 1e-10) to *Hodgkinia* and *Sulcia* references, respectively, using either qualimap (García-Alcalde et al. 2012) or bbmap (Bushnell 2014), and the mapped reads were used for final SPAdes assemblies. For *Hodgkinia* from *O. oregona*, PCR was used to close gaps and to verify rRNA operon sequences.

*Hodgkinia* and *Sulcia* genomes were annotated using a custom pipeline described previously (Łukasik et al. 2018), including curated sets of reference genes extracted from published genomes and tRNA annotation using tRNAscan-SE v.1.23 (Chan and Lowe 2019).

### DNA Coverage and Genome Abundance Analyses

Raw reads from the bacteriome metagenome sequencing libraries were inspected for quality using FastQC version 0.11.7 (https://www.bioinformatics.babraham.ac.uk/projects/fastqc/) and trimmed using Trim Galore! version 0.6.1 (https://www.bioinformatics.babraham.ac.uk/projects/trim_galore/) to remove Illumina adapters and low-quality bases (Phred scores <10) from read ends, retaining read pairs in which each read has a post-trimming length of at least 20 bp (Martin 2011).

For comparative analysis of *Hodgkinia* and *Sulcia* DNA coverage, BowTie 2 indexes were built from the *Hodgkinia* and *Sulcia* genomes of each cicada species using BowTie 2 version 2.4.1 (Langmead and Salzberg 2012). Trimmed DNA reads from each sample were aligned to the corresponding indexes. This and all subsequent DNA and RNA alignments were carried out in BowTie 2’s –very-sensitive mode, and this and all output alignment files were binary-compressed and sorted by chromosome position using SamTools version 1.12.0 (Li et al. 2009). MetaBAT adjusted coverage for contigs representing *Hodgkinia* and *Sulcia* was obtained using the jgi_summarize_bam_contig_depths function in MetaBAT2 (Kang et al. 2019). Using contig lengths and adjusted coverage values, reads representing *Hodgkinia* and *Sulcia* coverage were calculated and converted to percentages of total processed reads for each library.

To determine the relative abundance of each *Hodgkinia* genome in cicadas hosting multiple *Hodgkinia* lineages, trimmedDNA reads from each sample were aligned to the corresponding *Hodgkinia* genomes, and the coverage proportions were calculated exactly as in the comparison of *Hodgkinia* and *Sulcia* coverage, this time giving the proportion of *Hodgkinia* DNA coverage contributed by each *Hodgkinia* genome hosted by a given cicada.

### RNA Coverage Analyses and rRNA Sequence Removal

For visualization of strand-specific RNA-seq coverage of the *Hodgkinia* and *Sulcia* genomes, reads from the metatranscriptome sequencing libraries were quality checked and trimmed according to the same parameters as the DNA reads (see section ‘DNA Coverage and Genome Abundance Analyses’). Reads from each library were aligned separately to the appropriate *Hodgkinia* and *Sulcia* genomes. The RNA alignments were then separated according to the DNA strand from which the alignments originated. Briefly, reads which were the second in a pair and which aligned to the forward strand were written to a separate file (`samtools view -f 128 -F 16`). This was repeated for reads which were the first in a pair and aligned to the reverse strand (samtools view -f 80). These alignment files were combined (`samtools merge`), collectively representing RNA coverage of the minus strand of the corresponding genome(s). Alignments representing RNA coverage of the plus strands of these genomes were separated with a similar set of commands (`samtools view -f 144, samtools view -f 64 -F 16, samtools merge`).

From these alignment files, per-base coverage values were obtained for each strand using the `genomecov -d` command in BEDTools v.2.24.0 (Quinlan and Hall 2010). Stranded per-base coverage values, along with annotation information extracted from the GFF annotation files using custom Python scripts were used to generate coverage plots with Processing v.3.5.4 in Python mode. Custom Python and Processing scripts can be accessed from the following GitHub repository: https://github.com/noah-spencer/Supplement-for-Spencer 2023. For final coverage plots shown in Figure 3 and Supplementary Figure S1, these steps were repeated using alignment files randomly downsampled with SAMTools to achieve approximately 450X and 3500X coverage of *Hodgkinia* and *Sulcia* genomes of interest, respectively.

Since substantial rRNA sequence coverage was detected in some libraries, the processed reads were subject to bioinformatic rRNA depletion prior to determining transcript counts. Briefly, trimmed reads were mapped to all of the corresponding endosymbiont rRNA sequences. Mapped reads were removed from the resulting alignment files using SamTools (samtools view -f 4). Unmapped reads were converted back to paired-end FASTQ files using the SamToFastq function in Picard Toolkit v.2.23.7 (2020).

### Transcript Abundance and Antisense Transcription Analyses

The rRNA-depleted reads were aligned to the corresponding *Hodgkinia* or *Sulcia* genome(s). Transcript counts for *Hodgkinia* and *Sulcia* genes were obtained using FADU, a drop-in replacement for transcript quantification tools like htseq-count that uses partial counts and EM algorithms to more accurately assign reads derived from polycistronic transcripts (as produced by operons) and gene-dense coding regions (Chung et al. 2021). FADU was run in -s “reverse” mode to accurately quantify these stranded RNA-seq data. A second run in -s “yes” mode was performed to quantify transcription on the opposite strand relative to annotated open reading frames (i.e. to quantify antisense transcription). Percent antisense transcription for each biological replicate was estimated as the total number of counts output by FADU is -s “yes” mode divided by the summed counts from both runs. The percentages reported represent averages across the all biological replicates included for a given cicada species.

Counts output by FADU for putatively functional genes (excluding tRNA and rRNA genes) were converted to Transcripts Per Million (Wagner et al. 2012) using a custom Python script by Arkadiy Garber (https://github.com/Arkadiy-Garber/BagOfTricks/blob/main/count-to-tpm.py). Statistical analysis of these TPM expression data was performed in R v.4.0.4. Semi-partial correlation analysis of TPM expression data, relative gene dosage, and gene copy DNA abundance was performed using the R package ppcor v.1.1 (Kim 2015).

### Genome Sequence-Based Analyses

Sequences spanning 50bp upstream of start codons in (A) all protein-coding genes, (B) the 15 protein-coding genes with the highest average TPM, and (C) the 15 protein-coding genes wit the lowest average TPM were extracted from the *Hodgkinia* and *Sulcia* genomes from *T. ulnaria* and used to make six logo plots with WebLogo (Crookes et al. 2004).

Protein alignments of all copies of RpoD represented in our data, as well as RpoD from *Hodgkinia* in *Diceroprocta semicincta* and from the free-living alphaproteobacterium *Methylobacterium oxalidis* (retrieved from NCBI, protein accessions ACT34206 and GEP04622.1, respectively) were performed using MUSCLE algorithm (Edgar 2004) implemented through the M-Coffee web server (Moretti et al. 2007) and then visualized using NCBI’s Multiple Sequence Alignment Viewer v.1.2.0.

#### Data Availability

All data described here are available from the NCBI Umbrella BioProject PRJNA386376. All cicada bacteriome metatranscritpome and metagenome sequencing libraries were deposited in the SRA database under BioProject PRJNA923375. Newly-generated genome assemblies for endosymbionts of *D*. near *semicincta*, *T. ulnaria*, *T. limbata*, and *O. oregona*, as well as the corresponding SRA experiments (containing the raw reads), are available under BioProjects PRJNA923375, PRJNA512238, PRJNA246493, and PRJNA385844, respectively.

## Acknowledgements

We thank DeAnna Bublitz and Katherine Nazario for help with specimen collection and Arkadiy Garber for bioinformatics and programming assistance. This work was supported by the National Science Foundation [IOS-1553529 to J.P.M. and 026257-001 to N.J.S.]; the National Geographic Society [9760-15 to P.Ł.]; and the Gordon and Betty Moore Foundation [GBMF5602 to J.P.M.].

---

**Supplementary Table S1.**
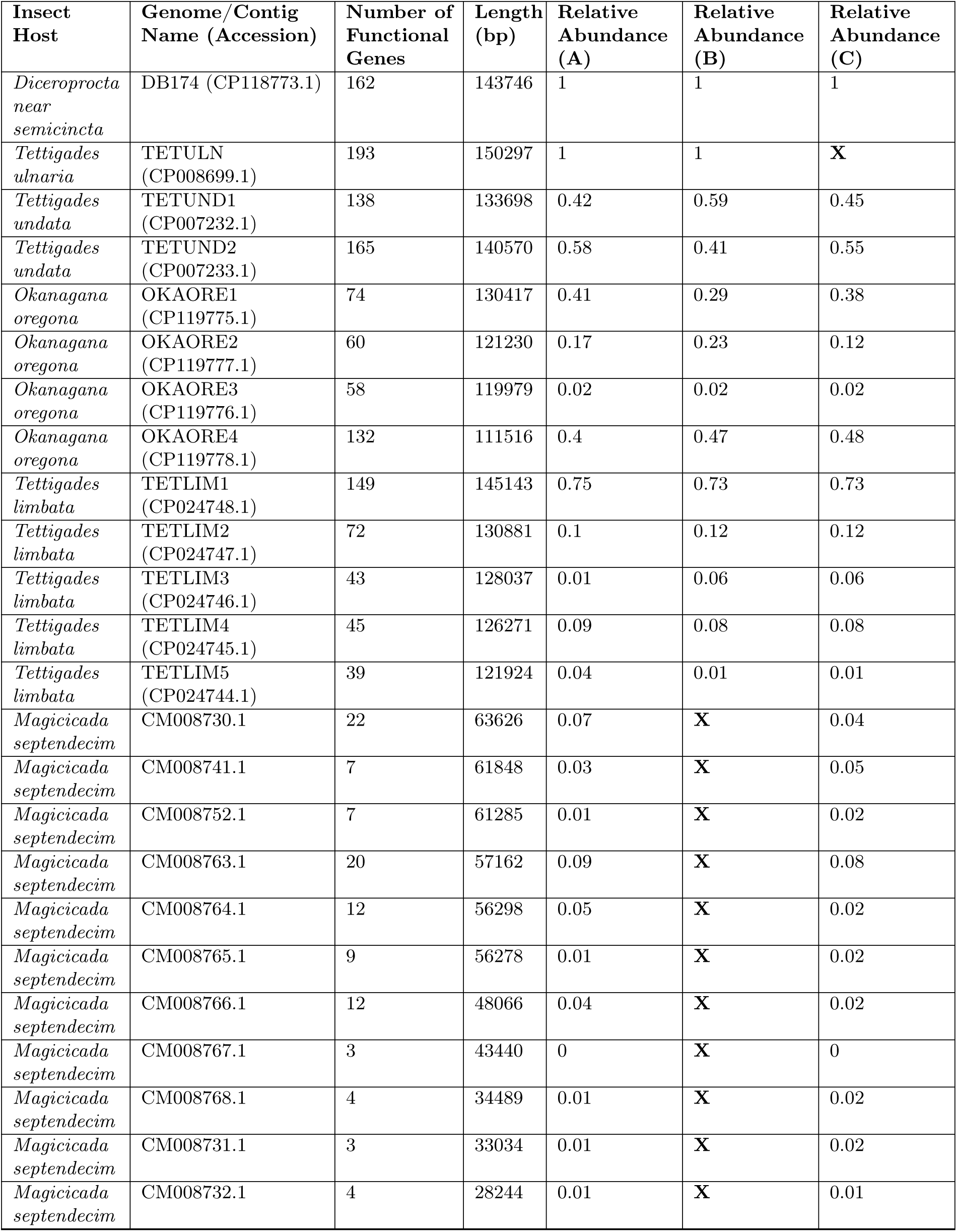
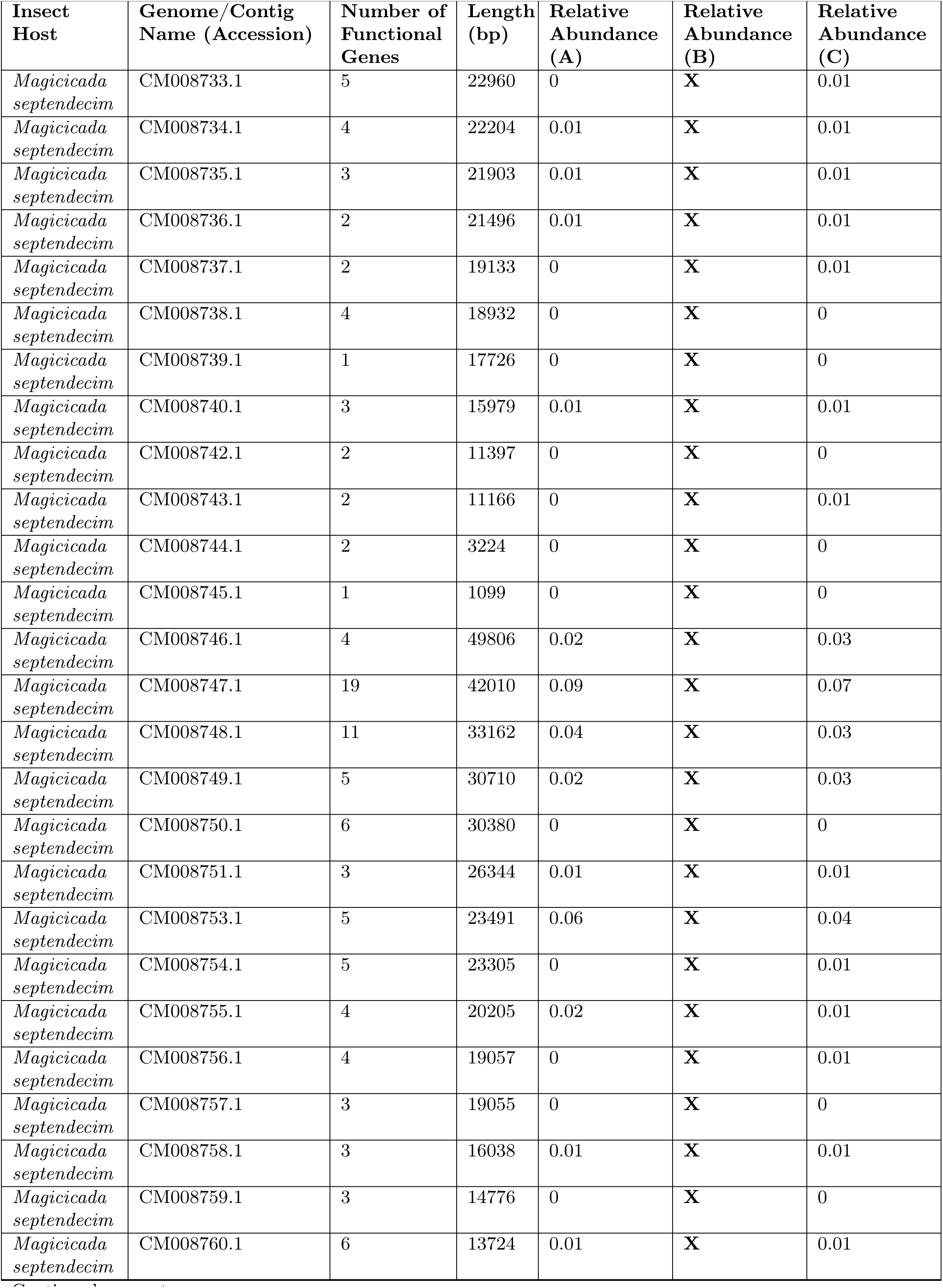
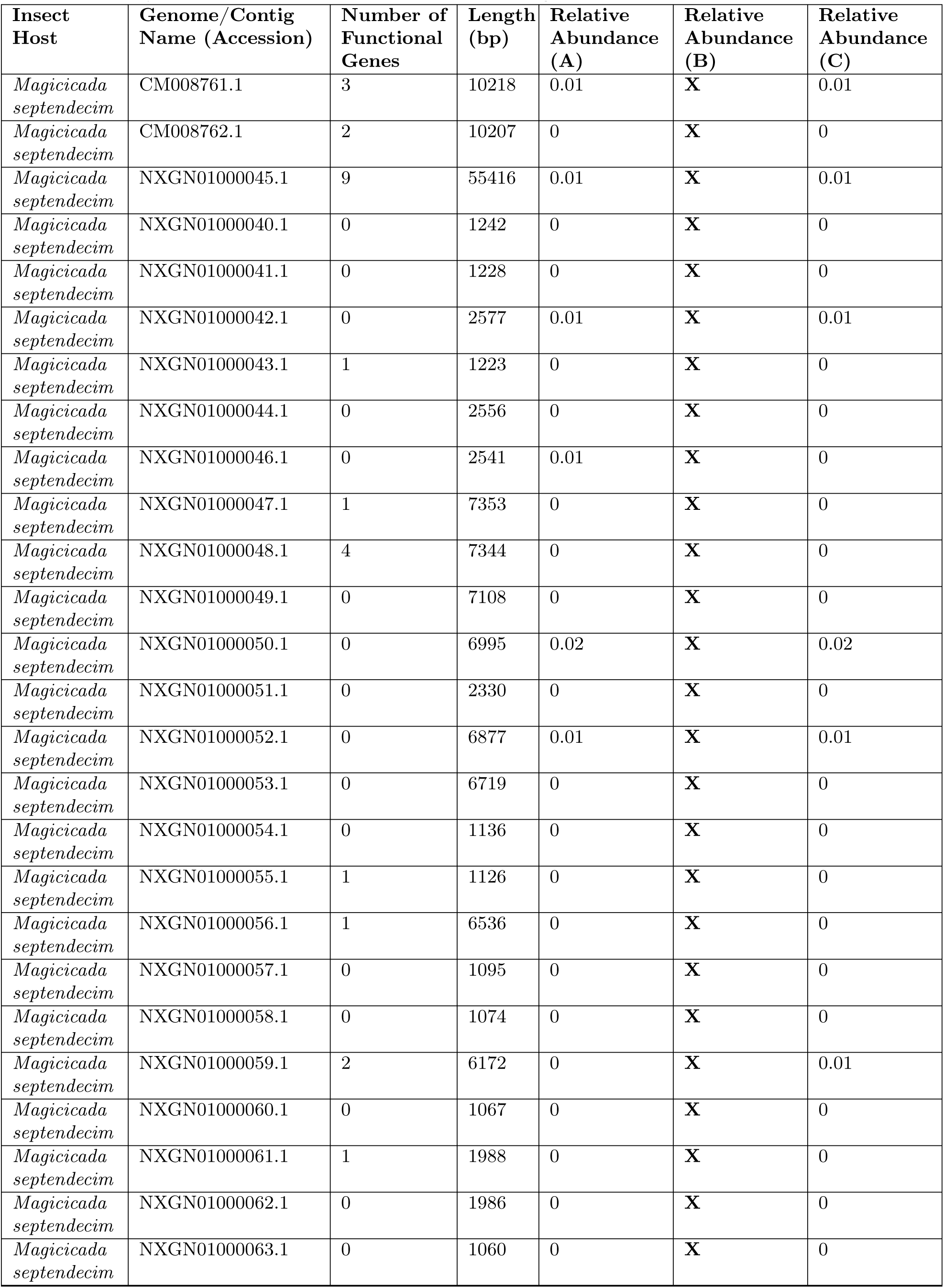
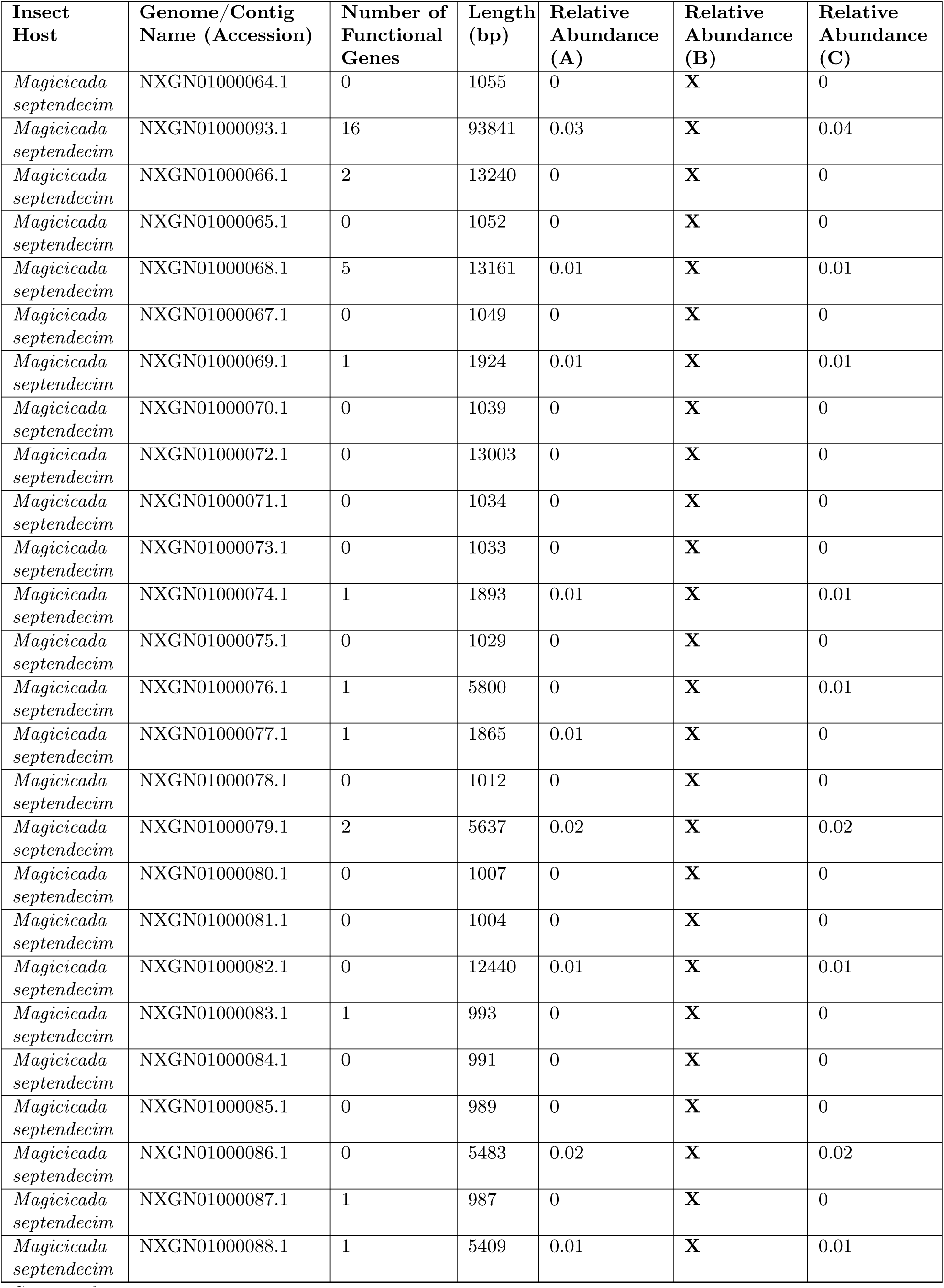
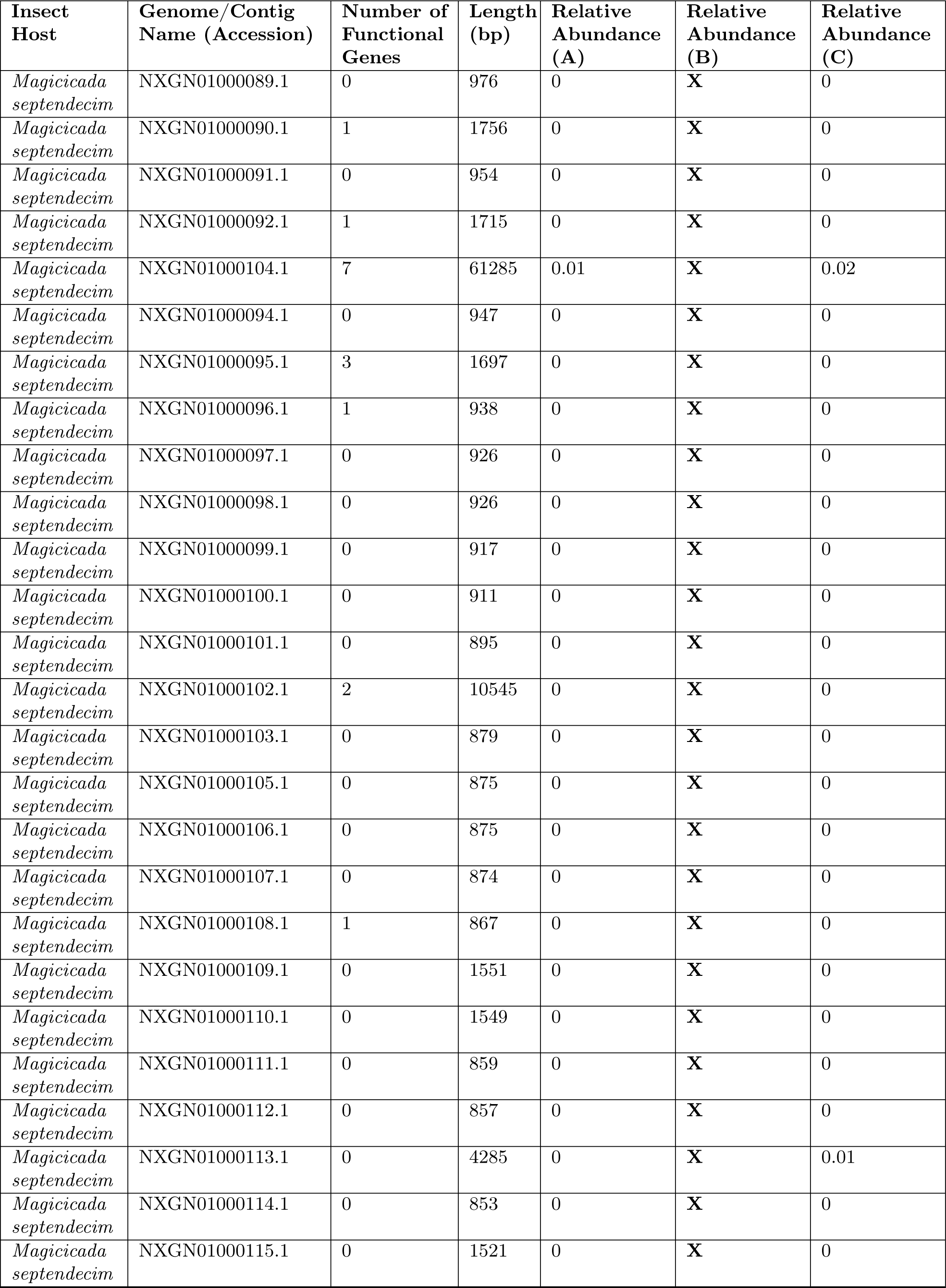
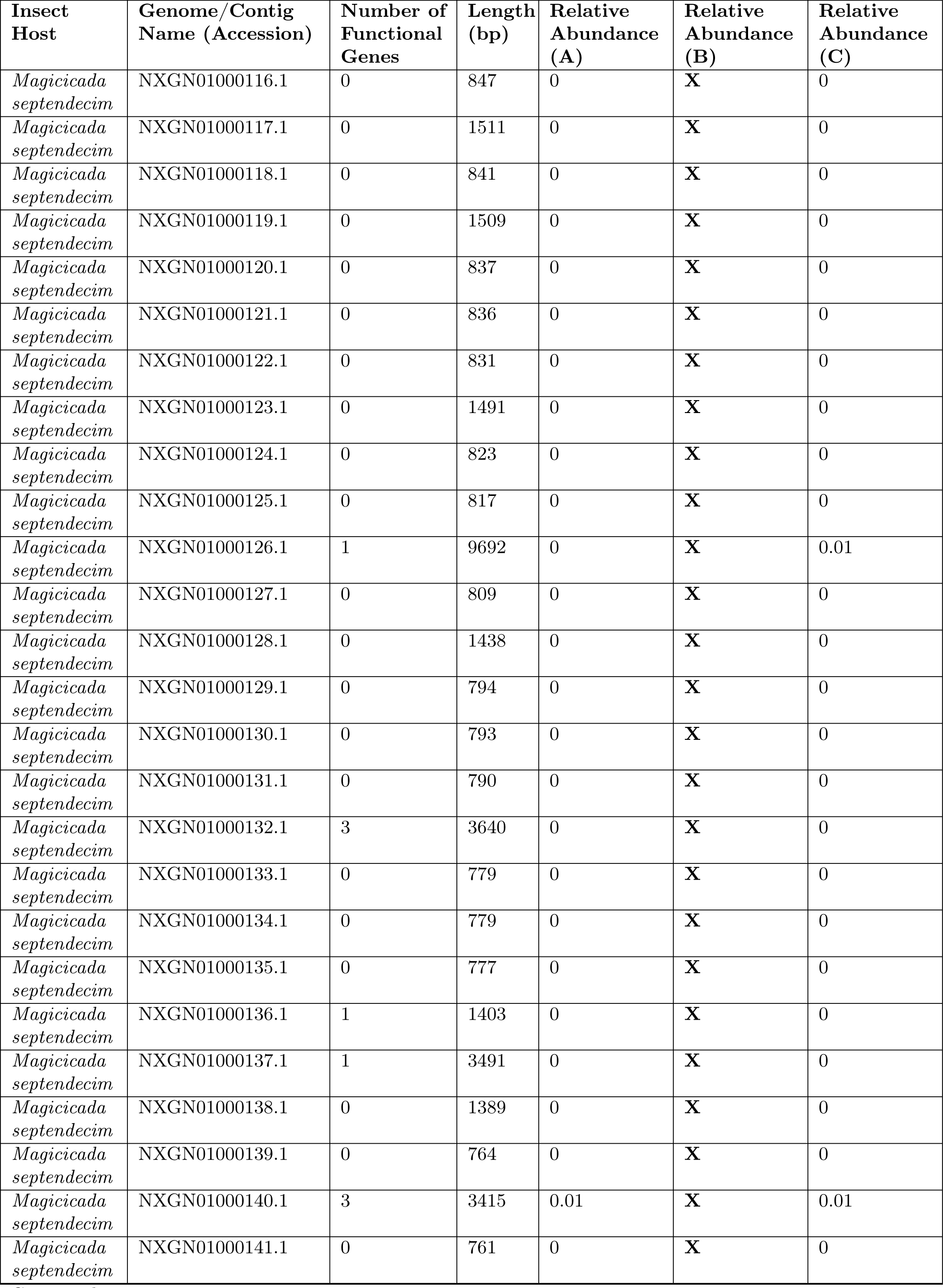
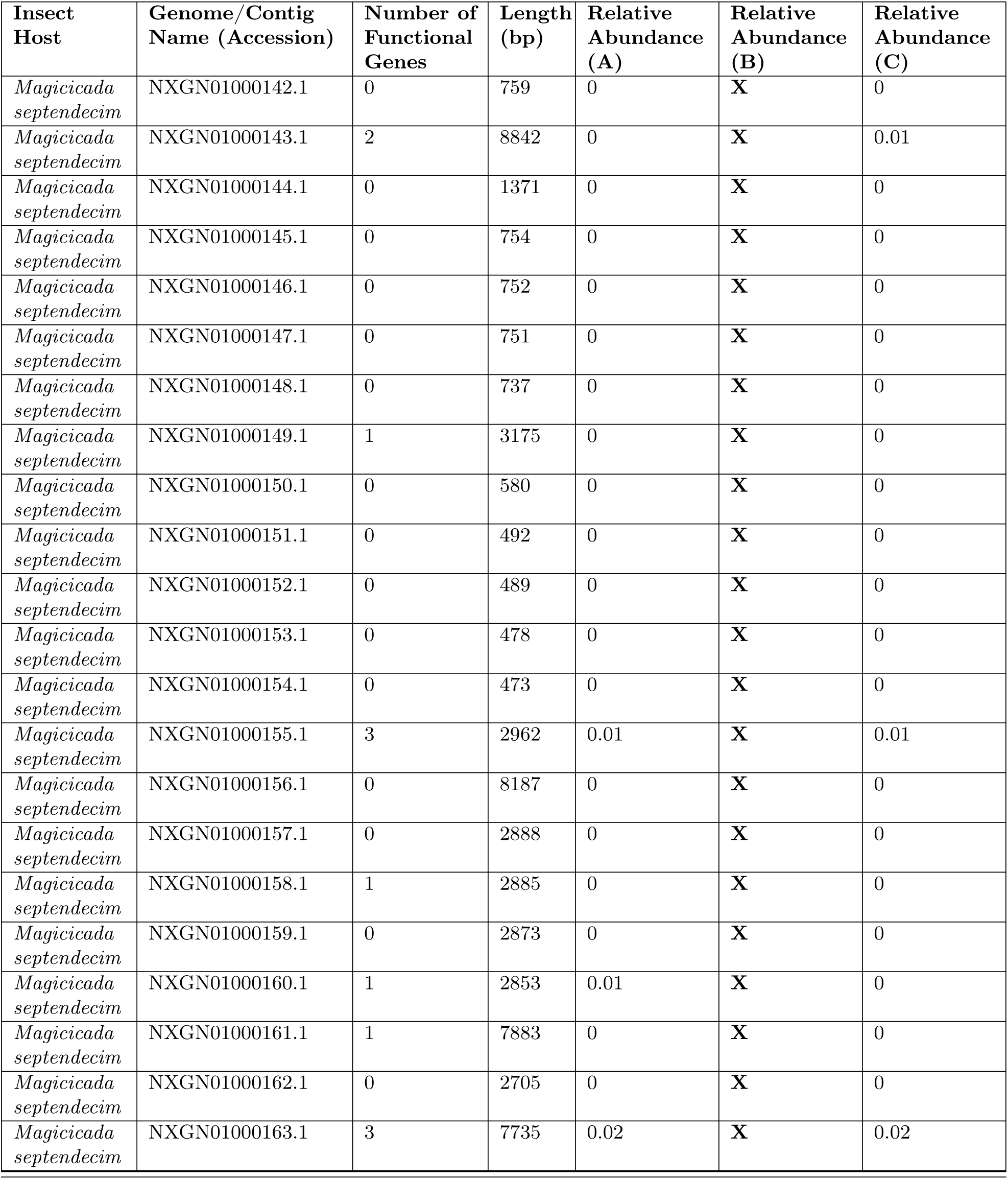
*Hodgkinia* genome complexity and abundance ratios from each specimen analyzed. Relative abundances are given for each biological replicate (labeled A, B, and C) of a given cicada species. A relative abundance value of "X" for any genome/sample combination indicates missing data.

**Supplementary Figure S1:**
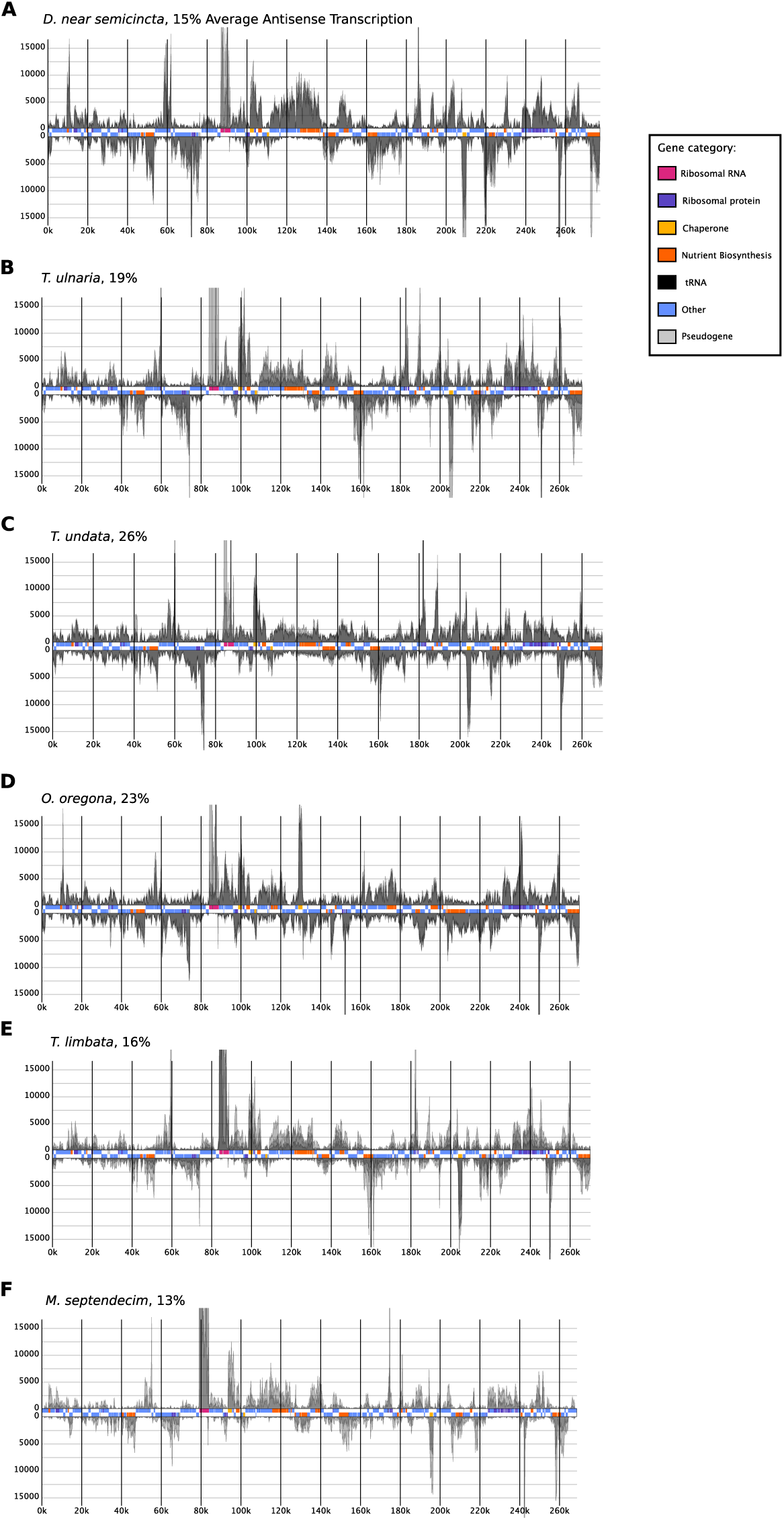
Sulcia RNA coverage plots and percent antisense transcription from all cicada species examined. Rectangles in the central track of each plot represent annotated genes and are colored according to functional categories. Positive and negative Y axes correspond to coverage of unfiltered RNA-seq reads derived from the plus and minus strands of each chromosome, respectively. Coverage represents alignments downsampled to approximately 3500X mean coverage of each genome and is cropped at approximately y=±20000. Antisense countsas a percentage of sense+antisense counts (averaged across biological replicates) for each Sulcia lineage are shown next to the name of the corresponding host species.

**Supplementary Figure S2:**
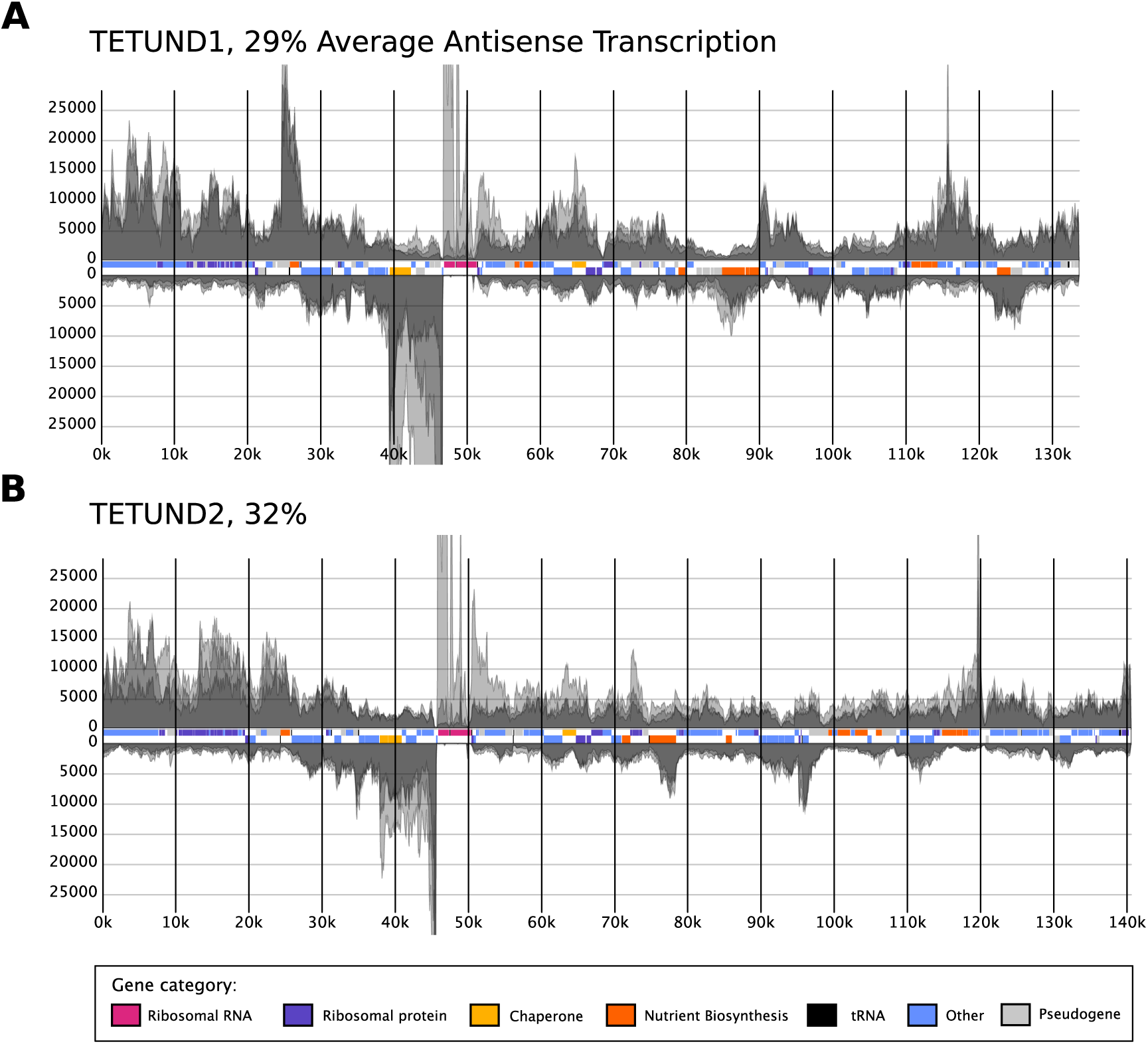
RNA coverage plots from all genomic lineages of Hodgkinia present in Tettigades undata. Rectangles in the central track of each plot represent annotated genes and are colored according to functional categories. Positive and negative Y axes correspond to coverage of unfiltered RNA-seq reads derived from the plus and minus strands of each chromosome, respectively and are cropped at approximately y=±30000. Antisense counts are shown as a percentage of sense+antisense counts (averaged across biological replicates) for each lineage.

**Supplementary Figure S3:**
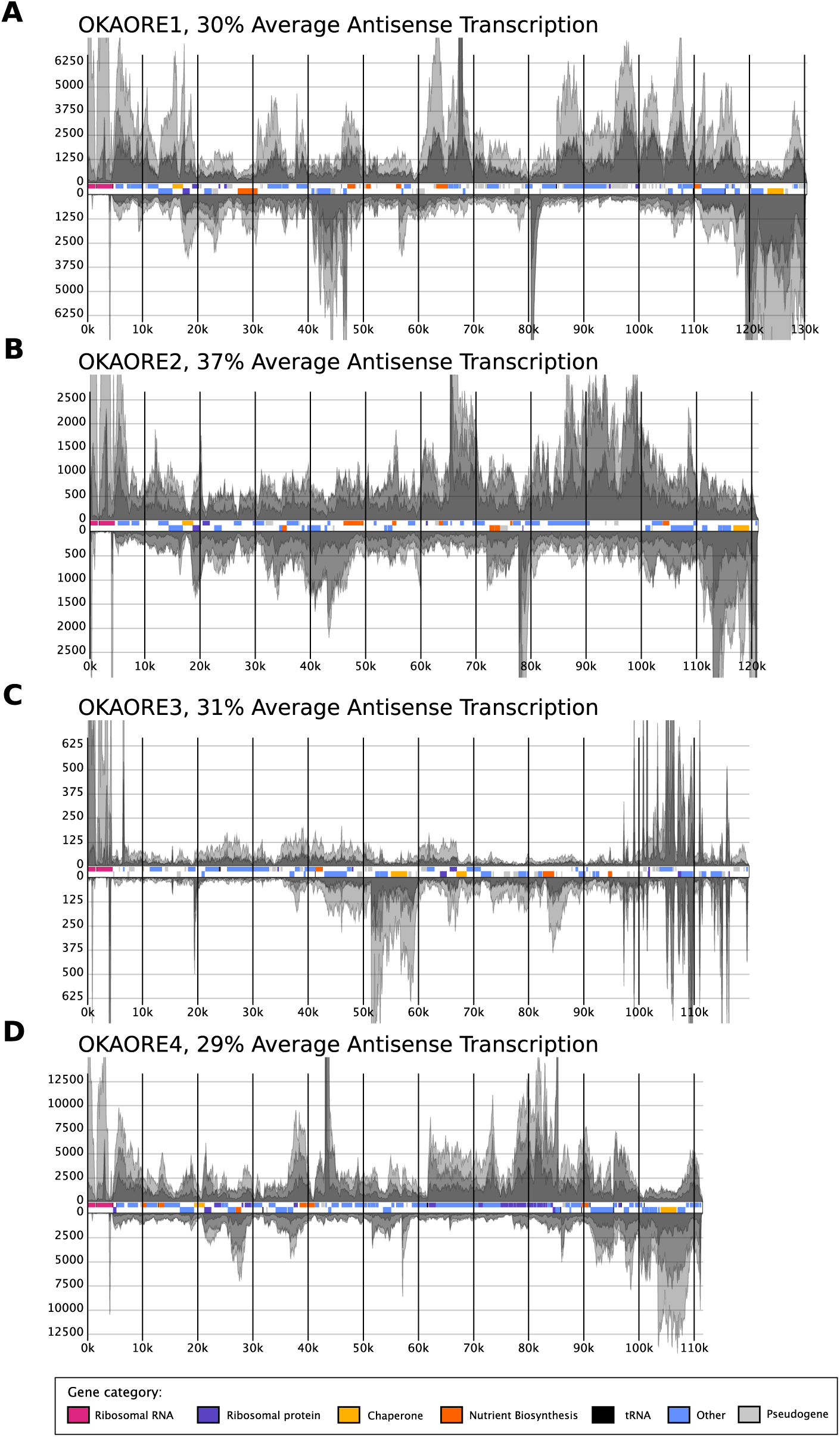
RNA coverage plots from all genomic lineages of Hodgkinia present in Okanagana oregona. Rectangles in the central track of each plot represent annotated genes and are colored according to functional categories. Positive and negative Y axes correspond to coverage of unfiltered RNA-seq reads derived from the plus and minus strands of each chromosome respectively and are cropped for readability. Antisense counts are shown as a percentage of sense+antisense counts (averaged across biological replicates) for each lineage.

**Supplementary Figure S4:**
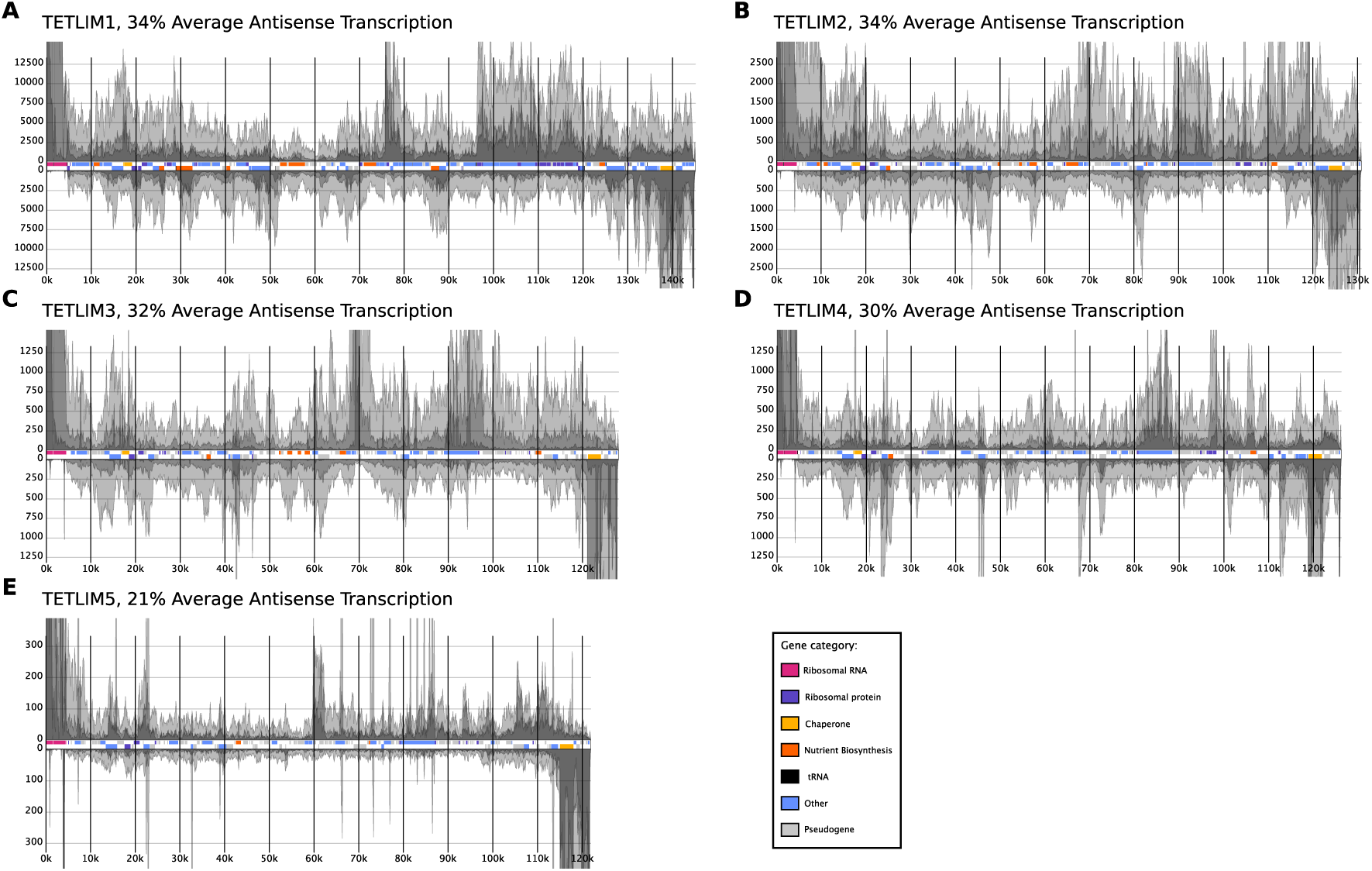
RNA coverage plots from all genomic lineages of Hodgkinia present in Tettigades limbata. Rectangles in the central track of each plot represent annotated genes and are colored according to functional categories. Positive and negative Y axes correspond to coverage of unfiltered RNA-seq reads derived from the plus and minus strands of each chromosome respectively and are cropped for readability. Antisense counts are shown as a percentage of sense+antisense counts (averaged across biological replicates) for each lineage.

**Supplementary Figure S5:**
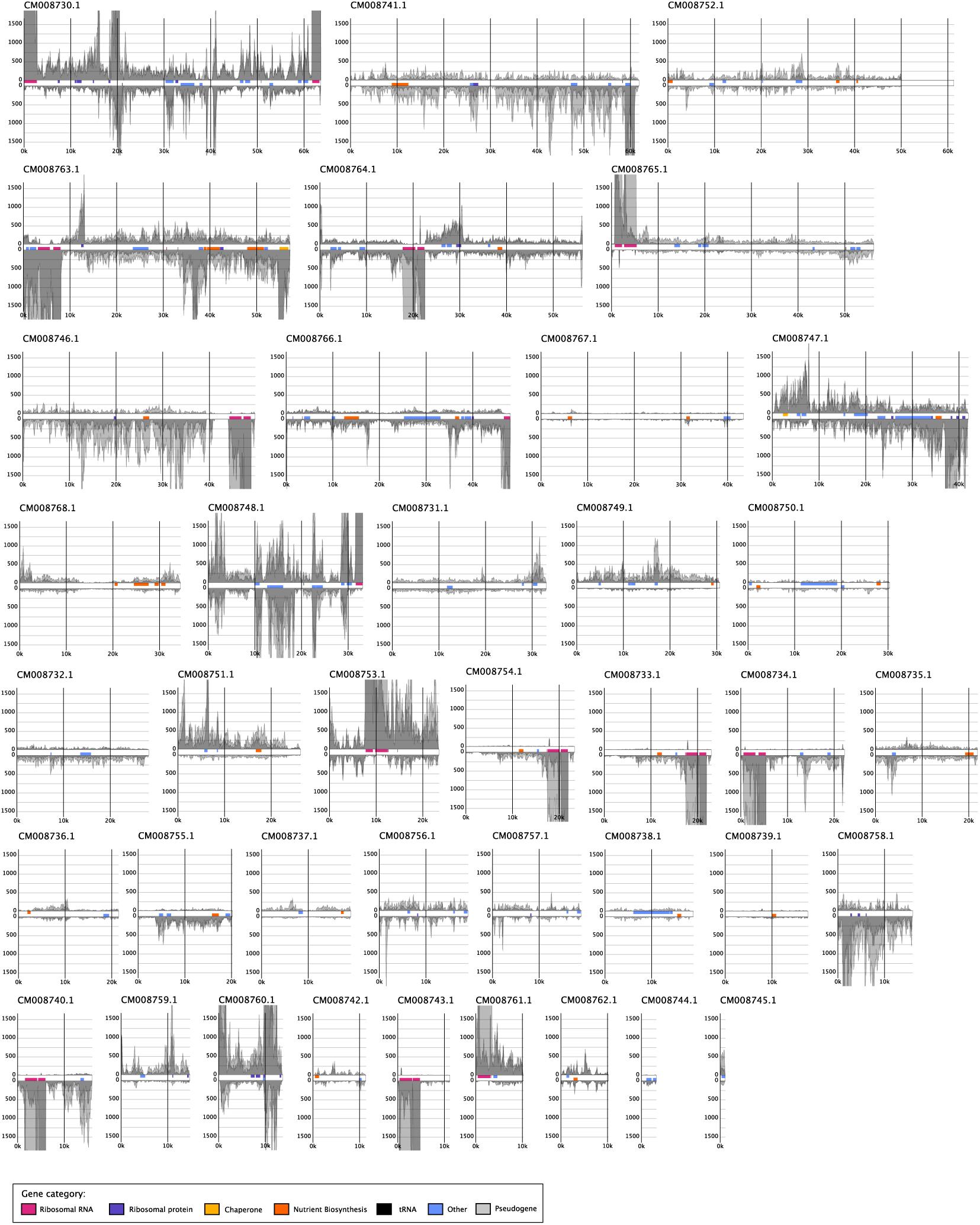
RNA coverage plots of all complete Hodgkinia genomic circles present in Magicicada septendecim. Rectangles in the central track of each plot represent annotated genes and are colored according to functional categories. Positive and negative Y axes correspond to coverage of unfiltered RNA-seq reads derived from the plus and minus strands of each chromosome respectively and are cropped for readability. Plots are ordered by contig length. On average, these Hodgkinia exhibited 26% antisense counts.

**Supplementary Figure S6:**
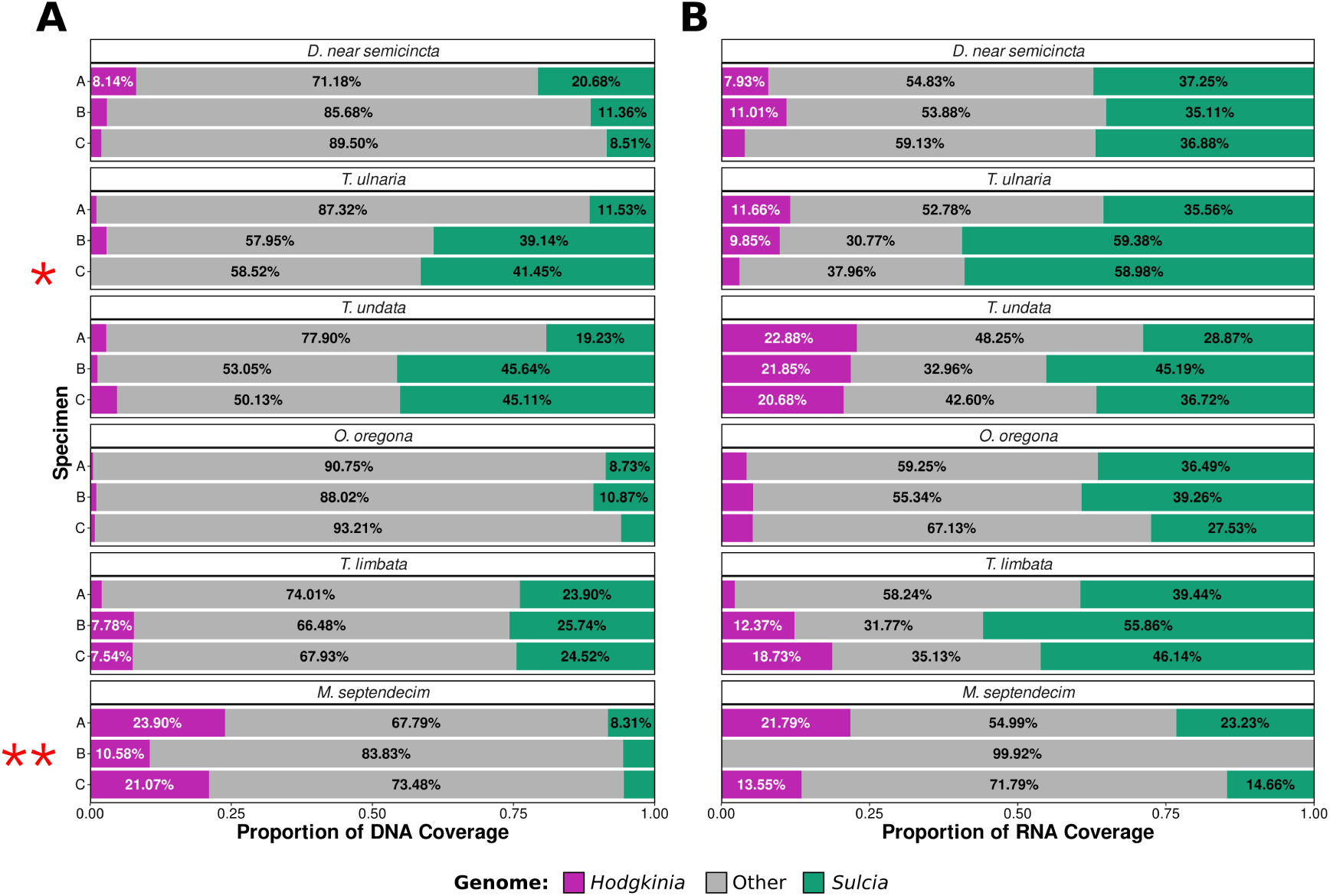
Contributions of Hodgkinia and Sulcia to DNA and RNA sequencing coverage prior to bioinformatic rRNA sequence removal. **(A)** DNA and **(B)** RNA sequencing coverage of Hodgkinia (magenta) and Sulcia (green) genomes as a percentage of total reads in triplicate biological replicates for each of the six cicada species examined. T. ulnaria specimen C (one asterisk) was excluded from Hodgkinia-based analyses. M. septendecim specimen B (two asterisks) was excluded from all further analyses.

**Supplementary Figure S7:**
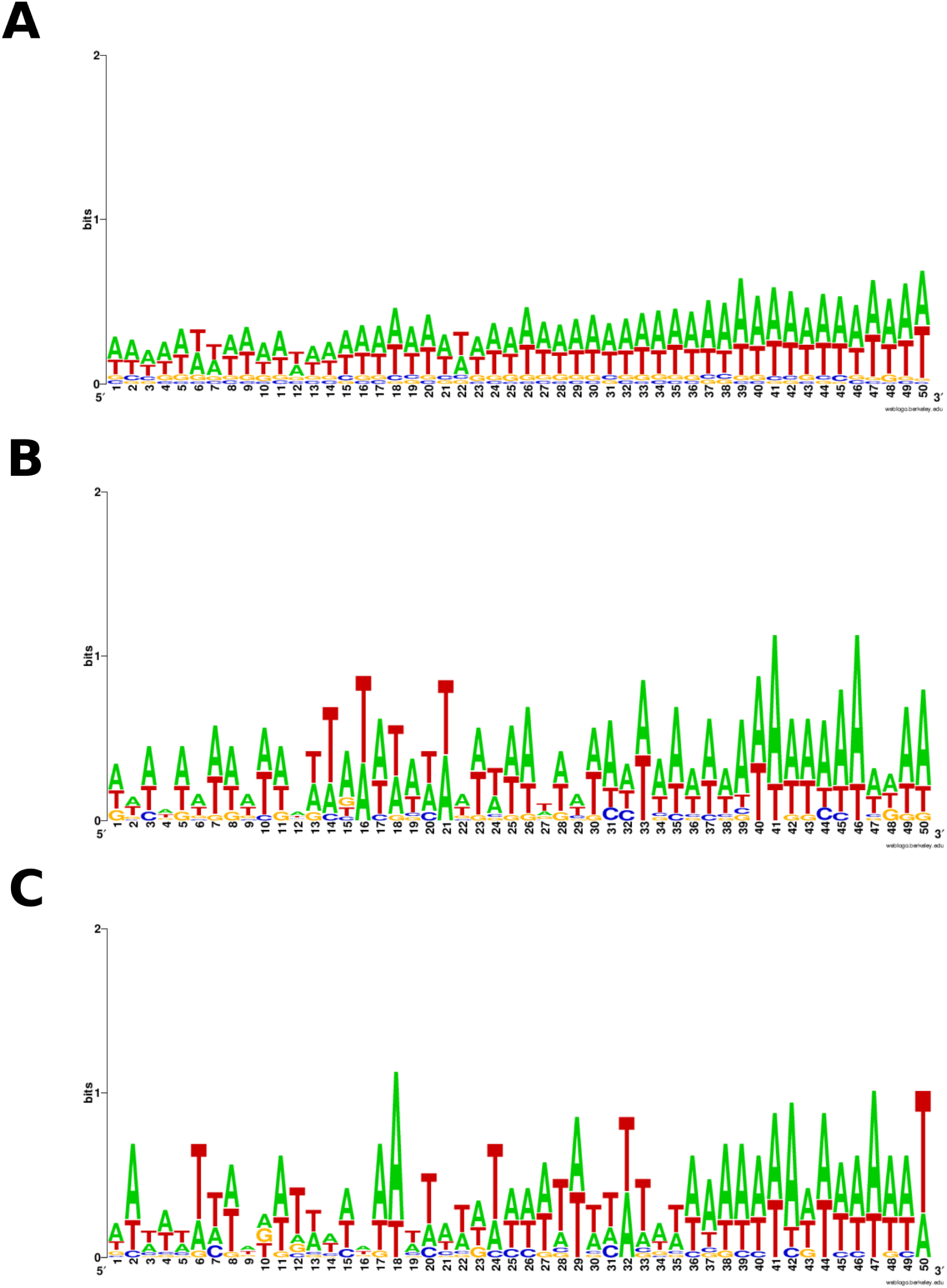
Logo plots of sequences immediately upstream from start codons in Sulcia from Tettigades ulnaria. Sequences were retrieved from either **(A)** all protein-coding genes, **(B)** the fifteen protein-coding genes with the highest average expression levels, or **(C)** the fifteen protein-coding genes with the lowest average expression levels. In each case, A and T bases are highly represented, which simply reflects the AT-rich nature of all Sulcia genomes.

**Supplementary Figure S8:**
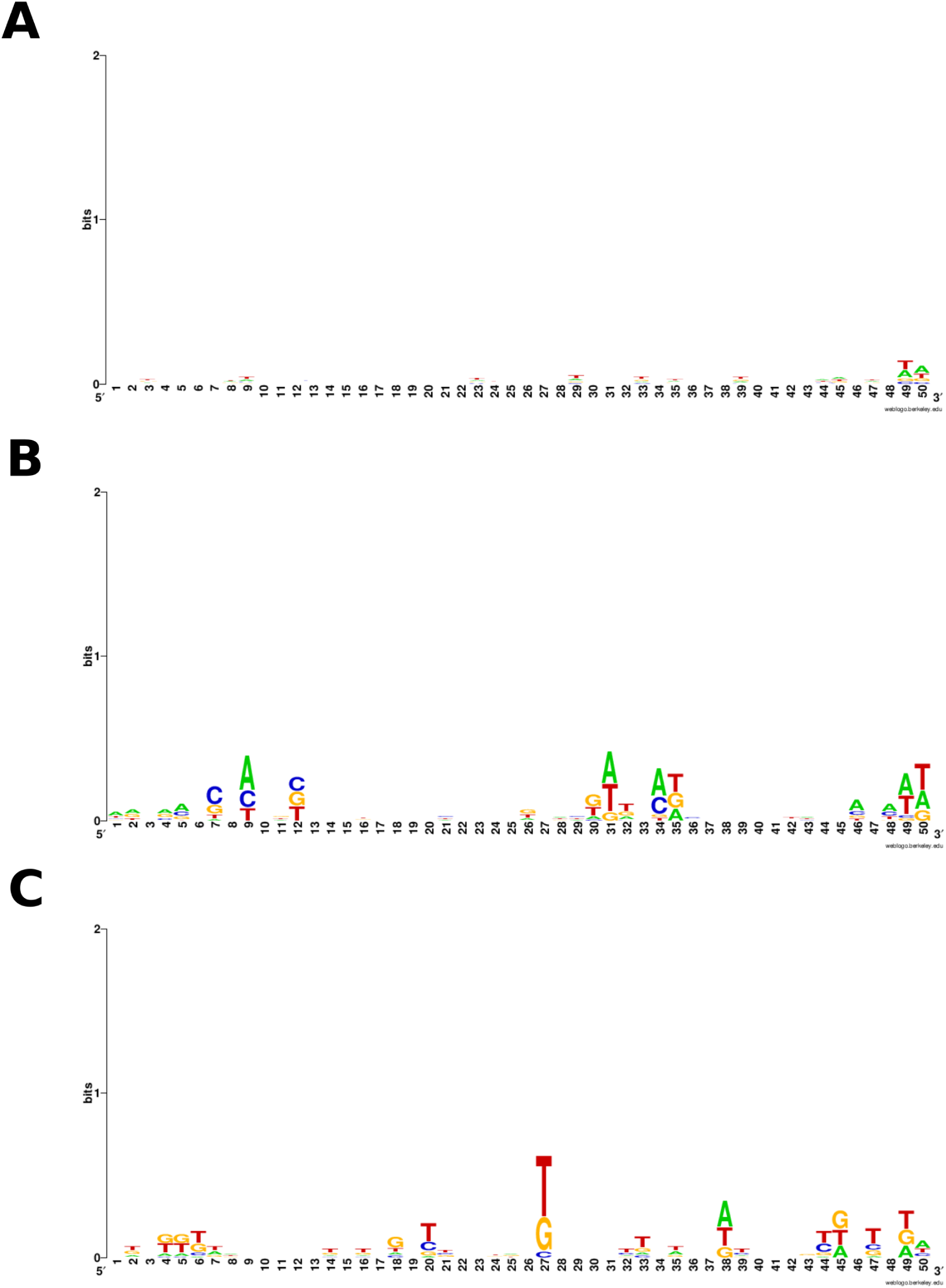
Logo plots of sequences immediately upstream from start codons in Hodgkinia from Tettigades ulnaria. Sequences were retrieved from either **(A)** all protein-coding genes, **(B)** the fifteen protein-coding genes with the highest average expression levels, or **(C)** the fifteen protein-coding genes with the lowest average expression levels.

**Supplementary Figure S9:**
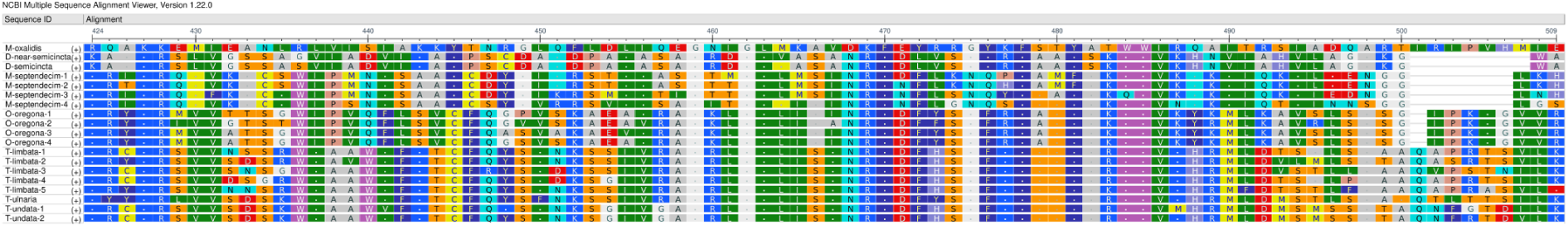
Protein alignment showing deletions in domain 3 of the σ70 factor (RpoD) in some Hodgkinia lineages. A portion of a protein alignment of all RpoD copies encoded by Hodgkinia lineages examined in this paper as well as RpoD from Hodgkinia in Diceroprocta semicincta and from the free-living alphaproteobacterium Methylobacterium oxalidis (retrieved from NCBI, protein accession: GEP04622.1) is shown. Hodgkinia from D. semicincta, D. near semicincta, and M. septendecim have gaps relative to other Hodgkinia lineages in the region spanning positions 501–507 of this amino acid alignment.

